# Multi-color live-cell STED nanoscopy of mitochondria with a gentle inner membrane stain

**DOI:** 10.1101/2022.05.09.491019

**Authors:** Tianyan Liu, Till Stephan, Peng Chen, Jingting Chen, Dietmar Riedel, Zhongtian Yang, Stefan Jakobs, Zhixing Chen

**Author notes:** Equal contribution (Tianyan Liu, Till Stephan). Corresponding author: Zhixing Chen.

## Abstract

Capturing mitochondria’s intricate and dynamic structure poses a daunting challenge for optical nanoscopy. Different labeling strategies have been demonstrated for live-cell stimulated emission depletion (STED) microscopy of mitochondria, but orthogonal strategies are yet to be established, and image acquisition has suffered either from photodamage to the organelles or from rapid photobleaching. Therefore, live-cell nanoscopy of mitochondria has been largely restricted to 2D single-color recordings of cancer cells. Here, by conjugation of cyclooctatetraene to a benzo-fused cyanine dye, we report a mitochondrial inner-membrane (IM) fluorescent marker, PK Mito Orange (PKMO), featuring efficient STED at 775 nm, strong photostability and markedly reduced phototoxicity. PKMO enables super-resolution recordings of inner-membrane dynamics for extended periods in immortalized mammalian cell lines, primary cells, and organoids. Photostability and reduced phototoxicity of PKMO open the door to live-cell 3D STED nanoscopy of mitochondria for three-dimensional analysis of the convoluted IM. PKMO is optically orthogonal with green and far-red markers allowing multiplexed recordings of mitochondria using commercial STED microscopes. Using multi-color STED, we demonstrate that imaging with PKMO can capture the sub-mitochondrial localization of proteins, or interactions of mitochondria with different cellular components, such as the ER or the cytoskeleton at sub-100 nm resolution. Thereby, this work offers a versatile tool for studying mitochondrial inner-membrane architecture and dynamics in a multiplexed manner.

## Introduction

Mitochondria are the powerhouses of the cell and govern key signaling pathways of cell homeostasis, proliferation, and death (1, 2). Due to their dynamic behavior, and abundant interactions with other organelles, mitochondrial research has been particularly driven by the development of fluorescence microscopy (3). However, the delicate double-membrane structure of mitochondria remains invisible using conventional fluorescence microscopes featuring a resolution limit of roughly 200 nm. Surrounded by a smooth outer membrane, the mitochondrial inner membrane (IM) forms numerous lamellar to tubular cristae, membrane invaginations which enhance the overall surface of the IM (4, 5). Crista junctions (CJs), small pores with a diameter of about 20 nm connect the invaginations to the residual part of the inner membrane and anchor the cristae along the organelle. In most cell types, cristae are densely stacked along the mitochondrial tubules, which can lead to crista-to-crista distances of way below 100 nm (6, 7). Due to this intricate arrangement of the cristae, transmission electron microscopy of fixed specimens remained the only tool to capture the unique mitochondrial membrane architecture for decades. However, coming to the era of super-resolution microscopy, stimulated emission depletion (STED) nanoscopy and structured illumination microscopy (SIM) have become the mainstay of live-cell mitochondrial imaging, with the former offering better spatial resolution of around 40-50 nm (3, 8–10) and the latter giving rise to faster imaging recording and longer imaging durations at about 100-120 nm resolution (11).

Like all nanoscopy techniques, STED nanoscopy relies on optimized fluorophores to reach its full potential. In the past several years, a handful of new mitochondrial labels have made possible the first live-cell nanoscopic captures of mitochondrial cristae and revealed their dynamic behavior (6, 12, 13). STED nanoscopy using MitoPB Yellow, Cox8A-SNAP-SiR, or MitoESq 635 all have showcased sub-100 nm resolution imaging of the IM (6, 12, 14). MitoPB Yellow (λ_ex_ = 488 mm, λ_STED_ = 660 nm) and MitoESq (λ_ex_ = 633 mm, λ_STED_ = 775 nm) are mitochondria-targeting, lipophilic dyes with remarkable photostability for time-lapse recordings. However, the widespread use of these two dyes has so far been prevented by phototoxicity or by the lack of combinability with other STED dyes. A different approach targeted the self-labeling SNAP-tag into the IM for subsequent labeling using the widely available silicone rhodamine dye (SiR, λ_ex_ = 633 nm, λ_STED_ = 775 nm) (15). Different to the membrane stains, this labeling strategy is generally applicable to the imaging of various mitochondrial proteins (6, 16–18), but it requires transfection or genetic manipulation and usually involves overexpression of the fusion proteins. STED nanoscopy of mitochondria labeled by SiR causes low photodamage (19) but suffers from rapid photobleaching during image acquisition, which strongly restricts the number of recordable frames when used to label cristae (6, 16).

From a technological point-of-view, the next challenge in nanoscopic live-cell imaging of mitochondrial physiology is to further expand the dimension of information, including long-term time-lapse tracking of mitochondrial dynamics, 3D analysis, and multiplexed imaging of various molecules inside and outside of mitochondria to unveil their interactions. To meet these demands, the next generation mitochondrial marker should feature: 1. Simple and robust protocol of highlighting mitochondrial structures in various cells and tissues; 2. High brightness and photostability, compatibility with a 775 nm STED laser which is available at most commercial STED microscopes, and compatibility with popular orthogonal nanoscopy dyes such as SiR for multi-color analysis; 3. Reduced phototoxicity to retain the integrity of mitochondria even under strong illumination.

As nanoscopy techniques generally require higher light doses to squeeze additional spatial and temporal information, photodamage and photobleaching can become a pronounced yet often under-evaluated technical hurdle for analyzing 4D physiology (19–21). We previously demonstrated that cyanine-cyclooctatetraene (COT) conjugates are gentle mitochondrial markers which allow extended time-lapse confocal and SIM imaging (22). Here, by adapting this class of labels for diffraction-unlimited STED nanoscopy we aim to provide a general and gentle tool that allows to embrace the era of 4D nanoscopy in mitochondrial imaging. We introduce PK Mito Orange (PKMO), an orange-emitting inner-membrane stain with minimal phototoxicity. PKMO is photostable and well-tailored for nearly all commercial STED microscopes. The cyclooctatetraene conjugated to PKMO depopulates its triplet state, markedly reducing the photodynamic damage during STED imaging. We demonstrate single-color time-lapse STED recordings of mitochondrial dynamics over the time course of several minutes and over 30 frames as well as 3D STED recordings of live mitochondria in cultivated cells. We demonstrate that PKMO can be combined with widely used fluorescent labels, enabling simultaneous localization of cristae along with mitochondrial protein complexes, mitochondrial DNA (mtDNA) or cellular organelles like the ER using multi-color nanoscopy. By resolving different cristae morphologies in living cells, we demonstrate that PKMO can pave the way for nanoscopy-based chemical and genetic screenings on mitochondria. Thereby, PKMO, in conjunction with other dyes for nanoscopy, will compose a palette which can complement classic transmission EM recordings used to investigate mitochondrial morphology and submitochondrial organization.

## Results

### PKMO is an orange-emitting cyanine-COT conjugate that stains the mitochondrial inner membrane

In our previous work, we introduced the two mitochondrial probes PK Mito Red (PKMR) and PK Mito Deep Red (PKMDR) which showcased low phototoxicity due to the use of the COT-conjugating strategy (22). PKMDR features an emission spectrum compatible with the more common 775-nm STED depletion, but the Cyanine 5 (Cy5) scaffold of PKMDR is prone to photobleaching compared to Cyanine 3 (Cy3) (23), giving only ~ 10 informative STED frames (*SI Appendix*, Fig. S1). In addition, PKMDR cannot be combined with popular live-cell compatible STED labels such as SiR due to overlapping emission spectra. To overcome this problem, we conjugated COT to Cy3.5 introducing the orange-emitting mitochondrial probe PKMO, which is photostable, well-suited for use with 775-nm STED lasers, and enables multicolor super-resolution (SR) imaging combined with green and far-red emitting probes. The chemical structures of PKMO and its analogue without COT, PKMO 0.9, are shown in Fig. 1a. Notably, these molecules feature a straightforward chemistry (< 7 steps from commercial material, see *SI Appendix* for detail), facilitating their large-scale production. The absorption and emission spectra of PKMO (Fig. 1b) demonstrate that the dye is tailored for STED nanoscopy using 561 nm excitation and a 775 nm depletion, a configuration implemented in most commercially available STED microscopes. Among the popular orange-red dyes, PKMO exhibited very good photostability in hydrophobic PMMA film (*SI Appendix*, Fig. S2). Moreover, PKMO led to significantly reduced singlet oxygen generation *in vitro* compared to PKMO 0.9 or TMRE as measured by the 1,3-diphenylisobenzofuran (DPBF) decay assay (24) under green LED illumination (50 mW/cm^2^ 520-530 nm) (Fig. 1c and *SI Appendix*, Fig. S2) (25). The absolute singlet oxygen quantum yield of PKMO was measured as 1.7 ± 0.1 *10^-3^, which is ~4 fold lower than that of PKMO 0.9 (6.7 ± 0.5*10^-3^). These data corroborate previous results with Cy3/Cy5 (22, 26), suggesting that phototoxicity and photostability are two related yet different properties of fluorophores in the excited state and hinting to a generally reduced phototoxicity of the COT-conjugate PK Mito dyes. To assess phototoxicity at the cellular level, we labeled live HeLa cells with 1 μM PKMO or 0.65 μM PKMO 0.9 (to achieve the same staining brightness, *SI Appendix*, Fig. S3) and illuminated the cells for different periods in a high-content imager before assessing apoptosis using a Calcein AM stain. The half-lethal light dose for cells stained with PKMO 0.9 was reached after ~ 10-min illumination, whereas for PKMO the dose was reached after 20-25 min, supporting the notion that the COT-conjugation significantly reduces cellular photodamage induced by long-term illumination.

**Figure 1.**
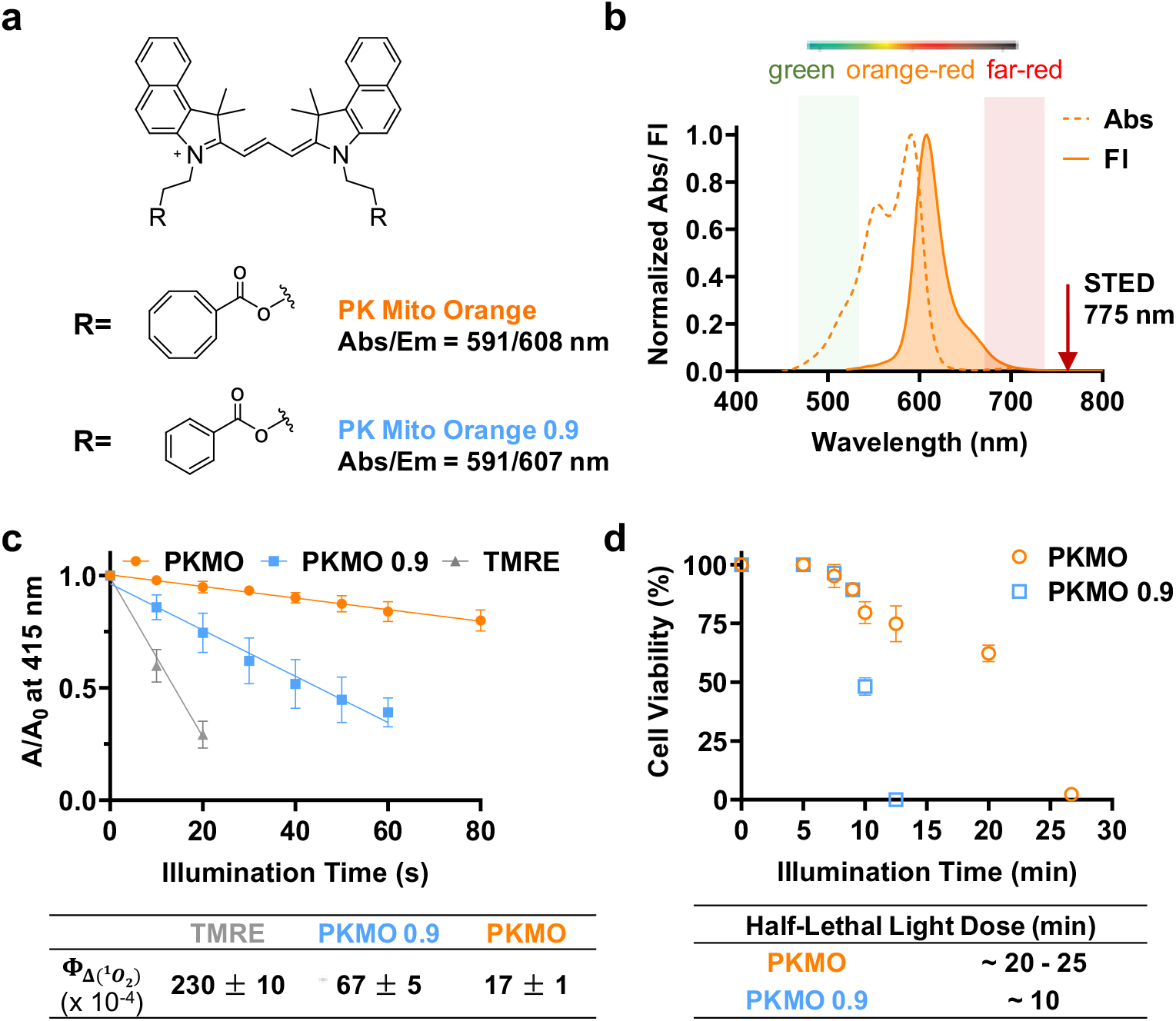
PK Mito Orange is a cyclooctatetraene-conjugated Cy3.5, featuring excellent photostability and remarkably low singlet oxygen generation. **a.** Chemical structures of PK Mito Orange and a benzoate-derived control compound. **b.** Absorption and fluorescence spectra of PK Mito Orange (PKMO) dwell in orange-red channel. The fluorescence can be potentially depleted using a 775-nm laser. **c.** Singlet oxygen quantum yields of PKMO and PKMO 0.9 measured using 1,3-diphenylisobenzofuran (DPBF) decay assay in acetonitrile. TMRE in MeOH (Φ_δ_= 0.012) was selected as a standard. **d.** Viability of PKMO (1 μM) and PKMO 0.9 (650 nM) stained HeLa cells after green LED light illumination (543-nm, 2.6 W/cm^2^). > 500 cells were examined in each time point of the three independent experiments.

### Long-term time-lapse recording and 3D-STED imaging of cristae with PKMO

To test the application of PKMO in super-resolution imaging, we stained COS-7 cells using 250 nM PKMO, resulting in brightly fluorescent mitochondrial network. PKMO showed excellent performance when recorded with a commercial STED nanoscope (Facility Line and STEDYCON, Abberior Instruments) equipped with 561 nm excitation and 775 nm depletion lasers, and like PKMR, the lipophilic and cationic PMKO specifically accumulated in the IM of mitochondria. We were able to record the mitochondrial networks across entire COS-7 cells and to capture individual mitochondrial cristae (Fig. 2a) at an optical resolution of down to 50 nm. (*SI Appendix*, Fig. S4). STED recordings revealed that mitochondria in COS-7 cells featured a highly ordered lamellar cristae architecture across the entire mitochondrial network (Fig. 2a and b). We measured the crista- to-crista distance from a selected area in Fig 2b and found distances of around 100 nm between closely spaced cristae (Fig 2c and *SI Appendix*, Fig S5).

**Figure 2.**
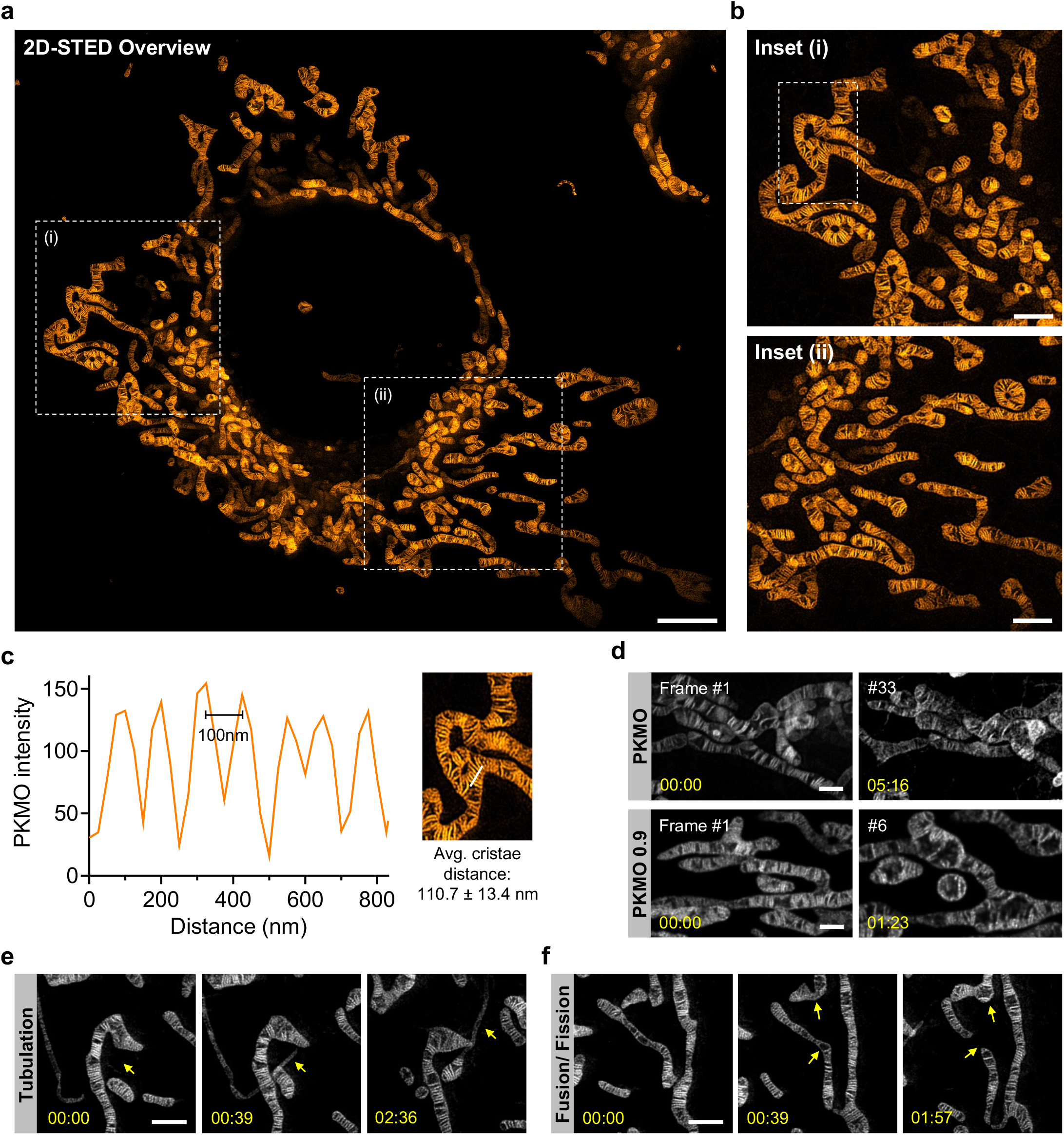
PKMO enables long time-lapse nanoscopic recording of mitochondrial cristae in COS-7 cells with minimal phototoxicity. **a.** 2D-STED recording of mitochondrial cristae of a COS-7 cell labeled with PKMO. Scale bar = 10 μm. **b.** Zoomed-in images of the white boxed areas (i, ii) in the **a**; the upper image is in (i), and the lower image is in (ii). **c.** Fluorescence intensity line profiles measured as indicated in the magnified view of white boxed area in the **b**. **d.** Time-lapse recordings of live COS-7 cell labeled with PKMO and PKMO 0.9; PKMO maintained both fluorescent signal and mitochondrial morphology during 30-frames of STED recording, while PKMO 0.9 caused visible mitochondrial swelling after 10 frames. **e.** Time-lapse STED nanoscopy recordings highlighting mitochondrial tubulation. Scale bar = 1 μm. **f.** Time-lapse STED nanoscopy recordings highlighting mitochondrial network dynamics such as fusion and fission. Scale bar = 1 μm. All the image data were deconvoluted and corrected for photobleaching.

In general, a limiting factor of live-cell STED imaging of mitochondria seems to be dye-induced phototoxicity since the fluorescence signals enriched in cristae membranes tend to decrease and can leak into the cytoplasm, accompanied by drop of mitochondrial membrane potential (MMP), when the cells are exposed to high light intensities (27). Indeed, COS-7 cells stained with PKMO 0.9 showed drastic mitochondrial swelling, accompanied by diffusion of the fluorescence signal after very few recorded frames (Fig. 2d). In contrast, the COT-conjugated counterpart, PKMO, enabled time-lapse STED recordings of cristae for 30 - 50 frames (Fig. 2d and *SI Appendix*, Fig. S6) before the onset of prohibitive photobleaching. The bottleneck of imaging PKMO in the STED mode was typically the eventual photobleaching, by which time the mitochondrial morphology still remained in shape. We conclude that because of reduced phototoxicity, mitochondria labeled by PKMO maintained a relatively healthy and intact morphology over the course of several minutes, showing mitochondrial dynamics including tubulation, fusion, and fissions (Fig. 2e, f).

The excellent 2D STED performance of PKMO encouraged us to test 3D STED nanoscopy of live COS-7 cells. Live-cell 3D STED nanoscopy is challenging due to several different limitations such as photobleaching of probes or rapid mobility of live mitochondria. Most importantly, phototoxicity effects accumulate by repeated scanning during Z-stacking, causing continuous swelling artifacts. Nevertheless, the gentle nature of PKMO allowed us to record the three-dimensional cristae architecture in mitochondria of a live COS-7 cell, at sub100-nm resolution using a 3D STED PSF. (Fig. 3a-b and *SI Appendix*, Fig. S7). Intriguingly, orthogonal cross sections revealed densely stacked and strongly titled cristae in the YZ section and hollow sections of mitochondria in the XZ section. Thereby, the data show the first three-dimensional mapping of cristae in live mitochondria, demonstrating that 3D STED can resolve complicated cristae arrangements, which would be barely visible using 2D nanoscopy.

**Figure 3.**
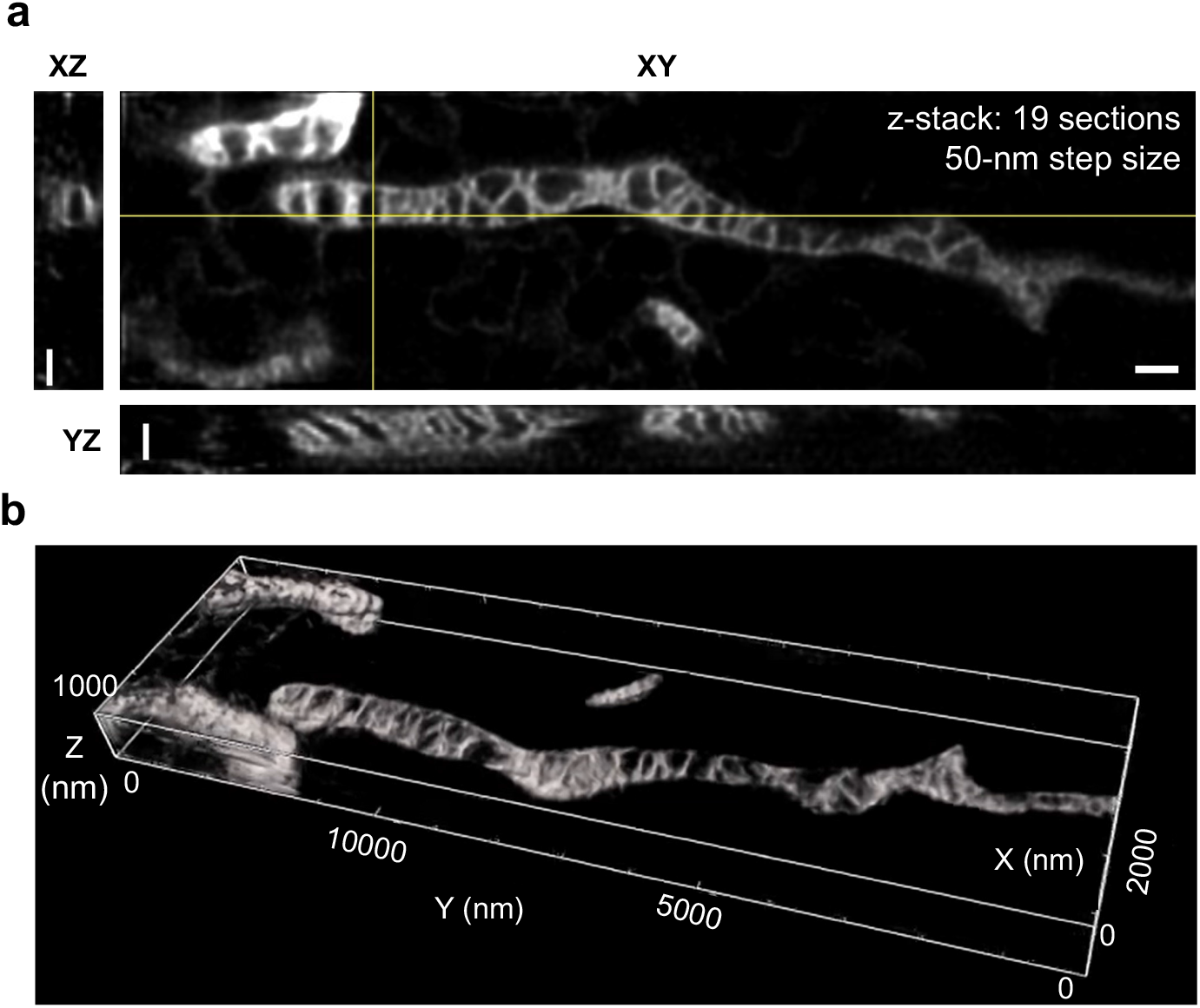
3D-STED imaging and reconstruction of cristae in COS-7 cells labeled with PKMO. **a.-b.** 3D live-cell STED recording of a mitochondrion from a COS-7 cell labeled with PKMO. **a.** Orthogonal cross-sections through 3D STED recordings. Scale bars: 500 nm. **b.** 3D reconstruction/ volume rendering of 3D STED data (Imaris).

### PKMO is a versatile cristae marker in various cell lines, primary cells, and organoids

In contrast to labeling strategies based on fluorescent proteins or self-labeling tags, PKMO spontaneously accumulates inside the IM of mitochondria. This behavior of the fluorophore should enable convenient mitochondrial labeling in various cell lines and tissues using a simple labeling protocol. So far, live-cell super-resolution microscopy of mitochondria has only been demonstrated in cultivated cancer cells, such as HeLa cells or COS-7 cells (6, 11, 12, 14, 16, 17, 22, 27–30), which feature relatively thick mitochondria and well-ordered lamellar cristae. We evaluated the performance of PKMO across other immortalized cell lines, namely HeLa cells and U-2 OS cells. As expected, both cell lines showed excellent PKMO staining and good STED performance (Fig. 4a, b). Cristae spacing in HeLa and U-2 OS cells was measured as 127 ± 33 nm (*SI Appendix*, Fig. S8) and 89 ± 22 nm (*SI Appendix*, Fig. S9), comparable with that in COS-7 cells (90 ± 24 nm, *SI Appendix*, Fig. S5). Moreover, STED recordings revealed that mitochondria in U-2 OS cells were generally more elongated and thinner than those in HeLa cells. The overall cristae arrangements appeared to be less regular compared to the striking lamellar cristae in COS-7 and HeLa cells (Fig. 2a and Fig. 4a).

**Figure 4.**
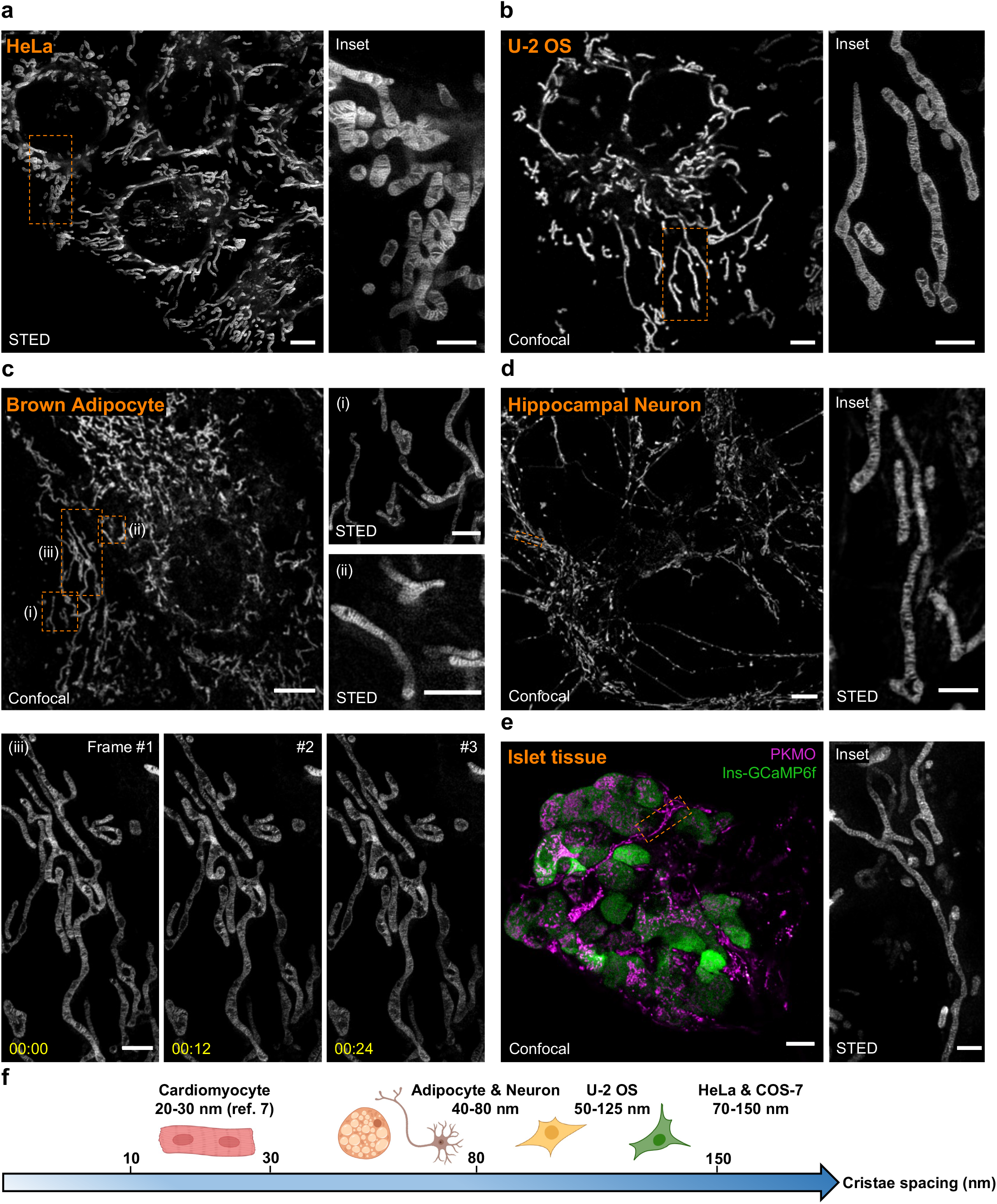
PKMO as a general mitochondrial cristae probe enables STED recordings on different cell lines, primary cells, and tissue. **a.** STED overview images (left) of live HeLa cells labeled with PKMO and zoom-in view (right) of the orange boxed area. Scale bar: overviews 5 μm, insets 2 μm. **b.** Confocal image of mitochondria labeled with PKMO in U-2 OS cell and zoom-in view of the dashed boxes in STED mode. Scale bar: overviews 5 μm, insets 2 μm. **c.** STED images of mitochondria labeled with PKMO in primary brown adipose cells (pBAcs). Confocal images of the mitochondria of pBAc and zoomed-in views or time-lapse recordings of the corresponding dashed boxes (i, ii, iii) in the overview. Scale bar: overviews 5 μm, insets 1 μm. **d.** Confocal image of mitochondria labeled with PKMO in primary hippocampal neurons (left) and zoom-in image of the dendrite in the boxed region is shown on the right. Scale bar: overviews 5 μm, insets 2 μm. **e.** Dual-color confocal image (left) of mitochondria (magenta, PKMO) and beta cells (green, Ins-GCaMP6f) in the primary islet tissue and zoomed-in STED images (right) of the corresponding orange boxed region (i, ii). Scale bar: overviews 5 μm, insets 1 μm. **f.** Cristae spacing ruler of various primary cells such as cardiomyocytes, adipocytes, and hippocampal neurons and cell lines such as U-2OS, HeLa, and COS-7. All STED data were deconvoluted and corrected for photobleaching.

Encouraged by its performance on immortal cell lines, we next aimed to test super-resolution mitochondrial imaging of primary cells and tissues labeled with our low-phototoxic probe PKMO (Fig. 4c-e). In primary brown adipocytes (pBAcs), PKMO delivered bright fluorescent labeling and STED nanoscopy recordings revealed a lamellar cristae architecture with dense cristae spacing of 63 ± 14 nm (*SI Appendix*, Fig. S10 and Table S1). Remarkably, we were also able to follow the dynamic behavior of cristae in pBAcs using time-lapse STED nanoscopy, suggesting that PKMO could be a promising tool to support the investigation of crista dynamics (Fig. 4c and *SI Appendix*, Fig. S10 and S11). We further tested PKMO in live mouse hippocampal neurons for mitochondrial labeling and cristae imaging (Fig. 4d and *SI Appendix*, Fig. S12). Mitochondria were clearly labeled in the cell bodies, axons, and dendrites (Fig. 4d, left panel), and STED imaging revealed dense cristae spacing at 62 ± 20 nm (*SI Appendix*, Fig. S13 and Table S1). Similarly, PKMO highlighted the fibrillar mitochondrial networks in rat cardiomyocytes (CMs) (*SI Appendix*, Fig. S14). However, STED nanoscopy was not able to visualize the cristae along the mitochondria of these cells. We attribute this to the intrinsic dense packing of cristae in CMs (~20 – 30 nm spacing according to electron microscopy (7) which is beyond the practical resolution of current commercial STED machines). Nonetheless, PKMO gave quantitative information on the cristae arrangement of various cells (Fig. 4f and *SI Appendix*, Table S1), circumventing time-consuming protocols of EM (13) and the potential sample distortion during chemical fixation (31). Strikingly, PKMO worked well also in primary tissues, as demonstrated on isolated mouse islet tissue (Fig. 4e). The confocal overview shown in Figure 4e (left panel) shows isolated islet tissue. Beta-cells expressing a genetically-encoded calcium sensor (Ins-GCaMP6f) are labeled in green, whereas the mitochondria of all cells within the islet tissue are labeled in magenta. Using 2D STED nanoscopy (Fig. 4e, right panel) we were able to capture the distinct lamellar cristae pattern in mitochondria from different cells within the islet tissue.

Together, these data demonstrate that PKMO allows a broad range of applications and can be used for convenient labeling of mitochondria across different cultivated cancer cells, isolated primary cells, and even to capture mitochondrial architecture in isolated tissues. We note, however, that super-resolution imaging of live primary cells remains challenging. First, many primary cells are often three-dimensional and small in size and have thinner and elongated mitochondria, which complicates super-resolution imaging. Second, labeling conditions need to be precisely adapted for different cells types. Third, primary cells are inherently fragile and tended to be more susceptible to photodamage compared to cancer cells. Especially in neurons mitochondria rapidly developed signs of significant and irreversible damage (*SI Appendix*, Fig. S12), which inevitably compressed the dynamic information to some extent. These challenges, in turn, underscore the importance of biocompatibility in the development of probes for nanoscopy (19–21).

### PKMO is compatible with various live-cell dyes and self-labeling tags for multiplexed recording

The double-membrane architecture of mitochondria creates different mitochondrial sub-compartments, which serve different purposes (Fig. 5a). Traditionally, biochemical analysis or electron microscopy have been used to analyze the submitochondrial localization of proteins. Multicolor nanoscopy has the potential to target the localization of biomolecules in live cells. PKMO is precisely designed for an application in multicolor nanoscopy due to its peak emission in the orange part of the spectrum. Excitation at 561 nm wavelength and depletion at 775 nm allows to combine PKMO with green fluorophores, which can be recorded in the confocal mode, or with a wide palette of deep-red fluorophores for dual-color STED nanoscopy. We tested the multi-color performance of PKMO in HeLa and COS-7 cells labeled with different transfection-free fluorescent probes or expressing fusion constructs with the self-labeling SNAP-tag or HaloTag (for an overview of labeling strategies, see Fig. 5a).

**Figure 5.**
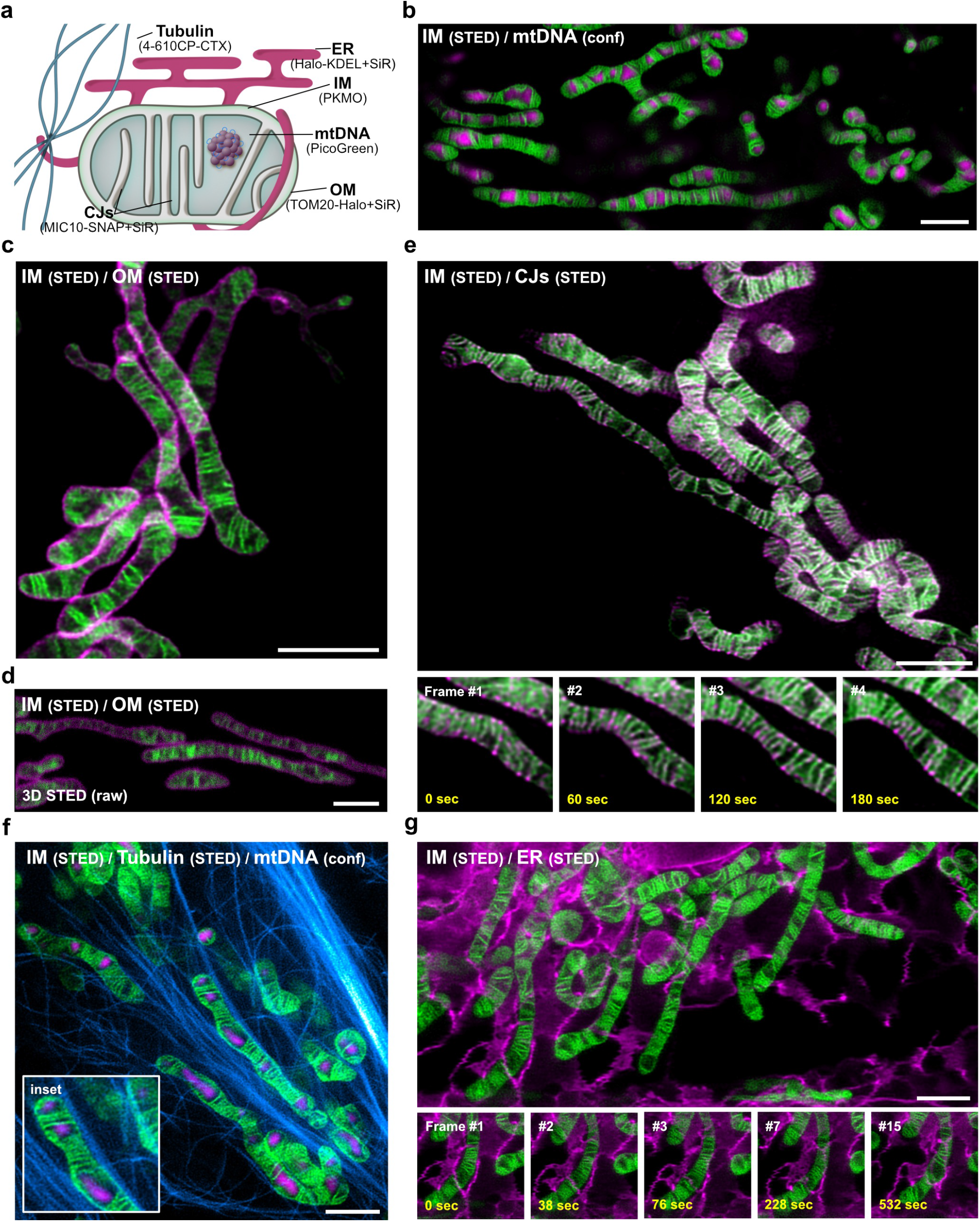
PKMO in conjunction with fluorogenic rhodamine probes enables nanoscopic mapping of mitochondria and the analysis of mito-organelle interactions using multi-color STED nanoscopy. **a.** Labeling strategies for multi-color live-cell recordings of mitochondria. Cartoon demonstrating the labeling strategies for mitochondrial sub-compartments and for mitochondria-interacting cellular structures. Abbreviations: ER (endoplasmic reticulum); IM (inner mitochondrial membrane); OM (outer mitochondrial membrane); CJs (crista junctions), mtDNA (mitochondrial DNA). **b-g.** Multi-color live-cell STED nanoscopy of HeLa and COS-7 cells labeled for the IM together with different mitochondrial targets or subcellular structures. IM was labeled using PKMO and recorded by live-cell STED nanoscopy (λ_ex_ = 561 nm, λ_STED_ = 775 nm). **b.** 2D single-color STED nanoscopy of mitochondria labeled for mtDNA. The mitochondrial DNA (mtDNA) was stained using PicoGreen and was recorded in the confocal mode (λ_ex_ = 488 nm). PKMO was recorded by 2D STED nanoscopy. **c-d.** Dual-color STED nanoscopy of mitochondria in COS-7 cells. OM marker TOM20-Halo was labeled using 647-SiR-CA (λ_ex_ = 640 nm, λ_STED_ = 775 nm). Cells were recorded by 2D STED in (**c**) or 3D STED nanoscopy (**d**). **e.** 2D dual-color time-lapse STED nanoscopy of mitochondria in HeLa cells. CJs were labeled by overexpression of MIC10-SNAP and staining with SNAP-Cell 647-SiR. Inset shows four consecutive frames illustrating cristae and CJ dynamics. **f.** 2D dual-color STED nanoscopy of mitochondria and microtubules in HeLa cells. Microtubules were labeled using 4-610CP-CTX (λ_ex_ = 640 nm, λ_STED_ = 775 nm). MtDNA was labeled using PicoGreen. Inset highlights contact sites of mitochondria and microtubules. **g.** 2D dual-color time-lapse STED nanoscopy of mitochondria and ER in HeLa cells. ER was labeled by overexpression of Halo-KDEL and staining with 647-SiR-CA. Inset highlights contact sites of a mitochondrion and ER over several time points. If not indicated otherwise, all image data were deconvoluted and corrected for photobleaching. Scale bars: 2 μm.

#### PKMO allows analyzing submitochondrial compartments

The mitochondrial DNA (mtDNA) is packed into nucleoids, which are located within the matrix of the organelle. In order to localize the mtDNA across the mitochondrial network of HeLa cells, we stained the cells with PKMO and the dsDNA-binding green-emitting probe PicoGreen. This allowed us to capture the spatial organization of the mtDNA (λ_ex_ 488nm, confocal) alongside the cristae (λ_ex_ 561 nm, λ_STED_ 775 nm) over the time course of several minutes (Fig. 5b and *SI Appendix*, Fig. S15). During this time, we observed significant remodeling of individual cristae and the overall mitochondrial network, but we found that the mtDNA remained trapped within the larger voids between groups of densely stacked cristae, suggesting that lamellar cristae generally act as diffusion barriers, which can separate different nucleoids (6).

Next, we aimed at a nanoscopic differentiation of the mitochondrial OM, IM and the CJs using dual-color live-cell STED nanoscopy. To this end, we expressed the OM protein TOM20 and the CJ marker MIC10 fused to the HaloTag and SNAP-tag, respectively. We stained the cells with PKMO and the deep-red fluorophores 647-SiR-CA or SNAP-Cell 647-SiR (15). Both PKMO (lex 561 nm) and SiR (lex 640 nm) are efficiently depleted using the 775 nm STED wavelength implemented in most STED microscopes. As expected, using dual-color live-cell STED nanoscopy, we were able to resolve the mitochondrial OM surrounding the PKMO-enriched IM using 2D STED as well as using 3D STED (Fig. 5c-d and *SI Appendix*, Fig. S16 and S17). Similarly, we were able to highlight the individual CJs alongside the cristae and to capture their dynamic movements over a few frames (Fig 5e and *SI Appendix*, Fig. S16). Thereby, the data demonstrate that combining PKMO with self-labeling tags and SiR, multi-color live-cell STED can help to analyze the sub-mitochondrial localization of different biomolecules, a task that is traditionally done using technically more demanding immunogold transmission electron microscopy (32). In contrast, STED manages to deliver similar information in living cells and does not require extensive sample preparation. Moreover, the approach allows drawing some information about the dynamics of the involved structures and biomolecules. However, we note that dual-color STED typically reduced the number of recordable frames compared to single-color recordings, which is mainly due to rapid photobleaching of SiR and occasionally because of enhanced photodamage (*SI Appendix*, Fig. S15-S16).

#### PKMO allows analyzing mito-organelle interactions

Mitochondria form a highly dynamic and interconnected tubular network that pervades the entire cytosol and features contact sites with different cellular structures. Especially important are interactions of mitochondria with the endoplasmic reticulum (ER). ER-mito contact sites are not only essential for the transport of lipids between both organelles (33, 34), but also for the overall dynamics of the mitochondrial network as they control fusion and fission of mitochondrial tubules (35–38). Importantly, ER-mito contacts have been considered increasingly important in the development of neurological disease in recent years (39). Similarly important for proper cellular function are interactions of mitochondria with the tubulin cytoskeleton, which determines the transport of the mitochondria (40). Therefore, an important aspect of future live-cell super-resolution microscopy studies will be high-resolution analysis of such interactions between mitochondria and cellular structures. To test the performance of PKMO in such scenarios, we labeled HeLa cells for the tubulin-cytoskeleton using the biocompatible fluorescent probe 4-610-CP-CTX (41) or for the ER by overexpressing Halo-KDEL (stained with 647-SiR-CA). Cells were co-labeled with PKMO and analyzed by live-cell dual-color STED nanoscopy. This approach allowed us to capture the tight spatial arrangement of mitochondria and the cytoskeleton at high spatial resolution (Fig. 5f). In addition, we could follow the interaction of the mitochondrial tubules with the ER over the time course of several minutes (Fig. 5g and *SI Appendix*, Fig. S18).

Taken together, live-cell STED nanoscopy of mitochondria labeled with PKMO allows rapid analysis of the sub-mitochondrial distribution of biomolecules and facilitates the recordings of dynamic interactions between mitochondria and other cellular structures.

### PKMO paves the way for a nanoscopic screening of the cristae architecture in living cells

The unique structure of the mitochondrial IM is inextricably linked to the functionality of mitochondria as the powerhouses of the cells. Defects of the cristae architecture cause malfunction of cellular respiration and are associated with neurodegenerative or cardiac diseases (42). Mutations or complete loss of proteins that control cristae morphology cause drastic remodeling of the cristae architecture (43–46). Traditionally, such changes of the cristae architecture have been analyzed using transmission electron microscopy of ultrathin sections of fixed cells. PKMO as a universal mitochondrial inner membrane marker that imposes little phototoxicity opens the door to perform such analysis also in living cells. As an example, we show cells that have been genetically modified using CRISPR-Cas9 to silence the genes of MIC10 or MIC60, core subunits of the mitochondrial contact site and cristae organizing system (MICOS) that controls CJ formation. We labeled these cells with PKMO and PicoGreen and recorded them using 2D live-cell STED. STED recordings clearly showed that all three cell lines feature distinct cristae phenotypes (Fig. 6a-c). Whereas wildtype HeLa cells showed highly ordered lamellar cristae (Fig. 6a), the cristae appeared as singe- or double-layered tubes that lined the mitochondrial interior in the absence of MIC10 (Fig. 6b). This phenotype is caused by a reduction in the number of CJs and by a defective segmentation of the IM into individual cristae (17). MIC60 depleted HeLa cells showed a different phenotype (Fig. 6c). The mitochondrial network of these cells was largely fragmented, and the large spherical mitochondria accumulated multiple layers of cristae, since MIC60depletion causes virtually a complete disruption of CJs (16, 17). Transmission EM recordings of the same cell lines (Fig. 6a-c) demonstrated that live-cell STED delivers similar information regarding the overall cristae arrangements like the electron microscopy recordings of resin-embedded samples. In addition, multi-color recordings allowed drawing additional information that are barely extractable from EM recordings. Our recordings showed that MICOS-depleted mitochondria do not only feature disturbed cristae, but also feature an aberrant distribution of the mtDNA, which can accumulate into large aggregates (Fig. 6b-c).

**Figure 6.**
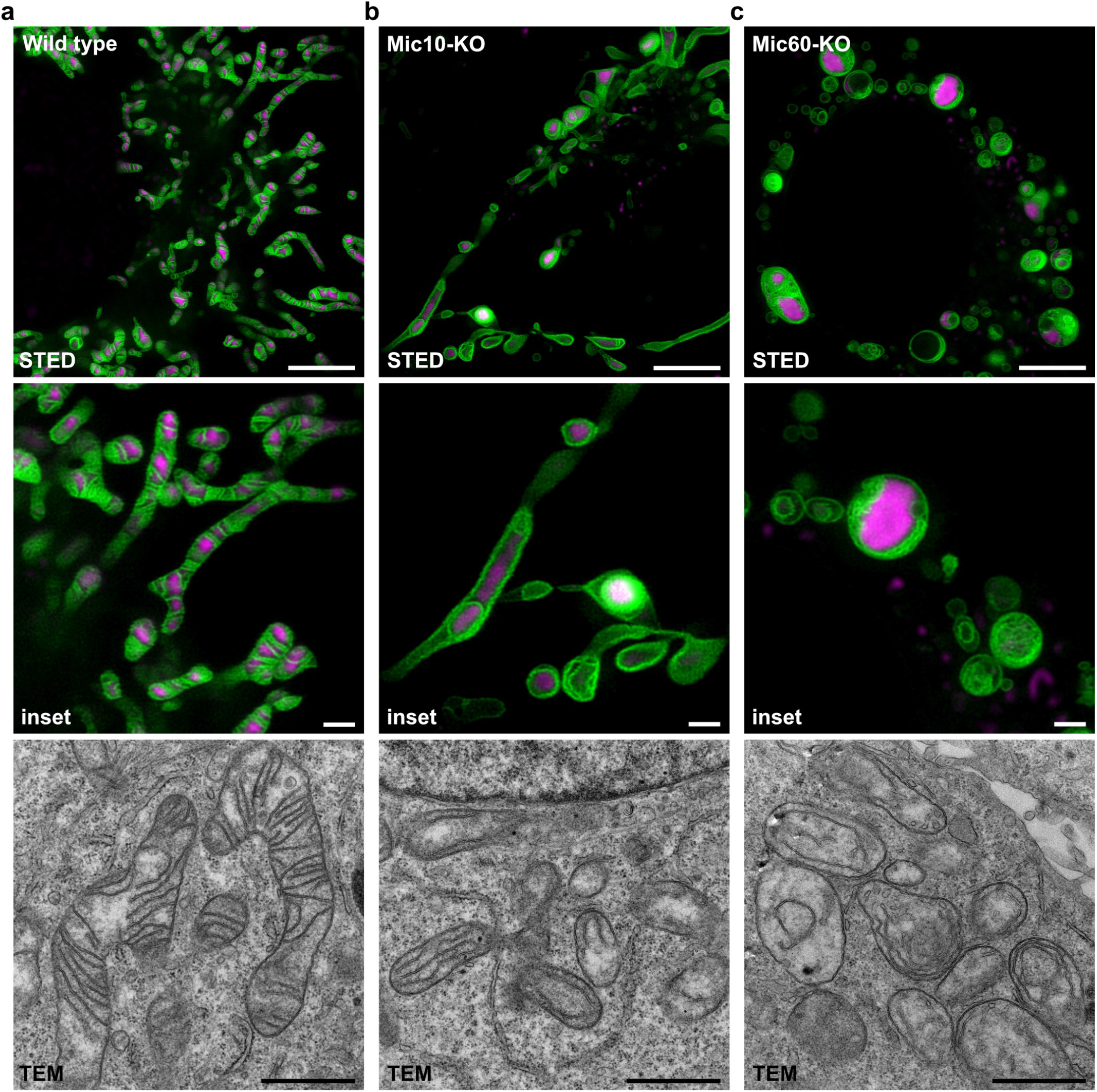
Analysis of cristae architecture using PKMO labeling and 2D live-cell STED nanoscopy. **a-c.** The mitochondrial inner membrane (IM) of HeLa cells was labeled using PKMO (λex = 561 nm, λ_STED_ = 775 nm). MtDNA was labeled using PicoGreen (λ_ex_ = 488 nm, confocal). Scale bars: overviews 5 μm, insets 2 μm. **a.** STED (upper panel) and transmission electron microscopy (TEM) recording (lower panel) of wild-type HeLa cells with typical lamellar cristae architecture (upper panel). **b.** STED and TEM recording of genome-edited HeLa cells lacking MIC10, a subunit of the mitochondrial contact site and cristae organizing system (MICOS complex). STED nanoscopy reveals tube-like and onion-like cristae arrangements and a disturbed arrangement of the mtDNA. **c.** STED and TEM recording of genome-edited HeLa cells lacking MIC60, the core subunit of MICOS. STED nanoscopy reveals a fragmented mitochondrial network, onion-like cristae arrangements, and aggregations of mtDNA. All STED data were deconvoluted. Scale bars: STED (overview) 5 μm, STED (insets) 1 μm, TEM 1 μm.

Our data demonstrate that PKMO can be used to analyze changes of the overall cristae architecture in live cultivated cells. If combined with automated imaging, this will open the door to a high throughput screening of compounds or siRNA libraries to conveniently analyze the impact of substances or proteins on the overall cristae morphology.

## Discussion and conclusion

Here we introduced PK Mito Orange (PKMO), a bright and versatile mitochondrial inner membrane stain with remarkably low phototoxicity that is tailored for multi-color STED imaging. From the perspective of probe development, the conceptual advance of this work lies in the introduction of triplet-state depleted dyes (47, 48) into STED nanoscopy to minimize photodamage. Phototoxicity has long been regarded as an important parameter in bioimaging practice and is often considered as a main hurdle of live-cell STED microscopy but approaches to chemically alleviate it were falling disproportionally short. This work showcased that a judiciously engineered mitochondrial stain can be gentle enough to allow ~20 z-stacks in a live cell for reconstructing cristae in 3D, opening future possibilities for the analysis of mitochondria. We thereby recapitulate the importance of assessing the phototoxicity of imaging probes, especially for nanoscopy(19, 20).

PKMO represents a timely addition to the toolbox of mitochondrial research using nanoscopic imaging. Its green excitation and orange-red emission profile are orthogonal with green dyes/FPs and the SiR-based far-red dyes, allowing unprecedented multiplexed nanoscopic recording of mitochondria-related processes in live cells. While GFP-based probes and sensors dominate diffraction-limited fluorescence microscopy techniques, the emerging red and far-red organic dyes, in conjunction with self-labeling protein tags, often outperform fluorescent proteins in nanoscopy due to superior optical properties (49, 50). The compatibility of PKMO with established dyes, the simple labeling protocol, the broad scope across different samples, and its compatibility with commercial STED microscopes promise to bring mitochondrial studies to the era of 3D nanoscopy. Crista dynamics, organelle interactions, and mitochondrial morphologies in genetically manipulated cells and even tissues can be examined in a multiplexed manner at unprecedented resolutions for long durations. Overall, this work opens exciting opportunities for mitochondria-related physiological and pathological research. We foresee that PKMO, as well as the other dyes of the mitochondrial palette in this work, will inspire additional methodologies ranging from probes, and algorithms, to instrumentations and will take part in various nanoscopic imaging practices in the era of 4D cellular physiology.

## Acknowledgements

This project was supported by funds from the Beijing Municipal Science & Technology Commission (Project: Z211100003321004 to Z.C.), Beijing Youth Top-Notch Talent Group (Project: 7350500012 to Z.C.), the Deutsche Forschungsgemeinschaft (DFG, German Research Foundation) through TRR 274/1 (2020) – ID408885537 (to S.J.), and the European Research Council through ERCAdG No. 835102 (to S.J.). We thank the Optofem Technology Limited for providing the instrument demonstrations of Abberior FACILITY and STEDYCON nanoscopes at Peking University. We thank Zhiwei He and Prof. Li Zhao for primary hippocampual neurons, Xuejiao Song and Prof. Xianhua Wang for adipocytes, Shiyan Tong and Prof. Liangyi Chen for islet tissues, Yingna Guo and Prof. Shiqiang Wang for cardiomyocytes, Junsheng Yang for helpful discussions on image processing, and Jan Keller-Findeisen for support with data analysis. We thank Gražvydas Lukinavičius for providing fluorescent probes and careful reading of the manuscript. We thank the analytical instrumentation center of Peking University and the NMR facility and optical imaging facility of the National Center for Protein Sciences at Peking University for assistance with data acquisition.

## Author contributions

Z.C. conceived the project and designed mitochondrial probes. Z.C., S.J., T.L., T.S. designed research. T.L., J.C and Z.Y. performed spectroscopy measurements, phototoxicity assays and STED microscopy of COS-7 cells, primary cells and tissues. T.S. performed STED imaging of HeLa and U-2 OS cells. P.C. performed chemical synthesis and characterizations. D.R. performed electron microscopy. T.L., T.S., S.J, and Z.C. analyzed data. T.L., T.S., and Z.C. wrote the paper with comments from all authors.

