## Supplemental Information for "Multi-color live-cell STED nanoscopy of mitochondria with a gentle inner membrane stain"

1) College of Future Technology, Institute of Molecular Medicine, National Biomedical Imaging Center, Beijing Key Laboratory of Cardiometabolic Molecular Medicine, Peking University, Beijing 100871, China
2) Peking-Tsinghua Center for Life Science, Academy for Advanced Interdisciplinary Studies, Peking University, Beijing 100871, China
3) Department of NanoBiophotonics, Max Planck Institute for Multidisciplinary Sciences, Göttingen 37077, Germany
4) Clinic of Neurology, University Medical Center Göttingen, Göttingen 37075, Germany
5) PKU-Nanjing Institute of Translational Medicine, Nanjing 211800, China
6) GenVivo Tech, Nanjing 211800, China
7) Laboratory of Electron Microscopy, Max Planck Institute for Multidisciplinary Sciences, Göttingen 37077, Germany
8) Fraunhofer Institute for Translational Medicine and Pharmacology ITMP, Translational Neuroinflammation and Automated Microscopy TNM, Göttingen 37075, Germany
9) Cluster of Excellence “Multiscale Bioimaging: from Molecular Machines to Networks of Excitable Cells” (MBExC), University of Göttingen, Göttingen 37099, Germany
10) Equal contribution

**This PDF file includes:**

Supplementary information text
- Material and Methods
- Figures S1 to S17
- Table S1

Chemical synthesis and characterization of new compounds

SI References

### Supplementary Information Text

#### Material and Methods

**UV-Vis and fluorescence spectroscopy**
Stock solutions of PKMO and PKMO 0.9 were prepared in DMSO solvent and diluted with methanol (MeOH) to 1 µM. UV-vis absorption spectra of sample solutions were measured using a Shimadzu UV3600Plus UV-VIS-NIR Spectrophotometer (Shimadzu, Japan) in a 1cm square quartz cuvette. Fluorescence emission spectra were measured using a Shimadzu RF-5301PC spectrofluorometer (Shimadzu, Japan) in a 1cm square quartz cuvette. The concentration of dyes was determined based on the absorbance at 591 nm ($\varepsilon_{MeOH} = 1.05\times10^5\ M^{-1}cm^{-1}$) (1).

**Quantum yield determination**
Stock solutions of PKMO and PKMO 0.9 were prepared in DMSO solvent and diluted with methanol or toluene to 1 µM. Absolute quantum yields of dyes were measured using PLQY (77-500K) (OXFORD microstat/FLS980, England) at room temperature. This instrument uses an integrating sphere to determine photons absorbed and emitted by a sample. Measurements were carried out using diluted samples (A < 0.1) at their excitation wavelength (560 nm).

**Bulk bleaching measurement**
Dyes were immobilized within polyvinyl alcohol (PVA) and polymethylmethacrylate (PMMA) films prepared using a spin coater (Setcas, SW-4A spin coater, China) and round-glass coverslip (diameter = 25 mm) as substrates. For PVA films, 100 µL 5% PVA (Yuanye, S27770-500g) in Mili-Q water containing 2 µM dyes were applied to the coverslip followed by spin coating (80,000 rpm 10 sec; 300,000 rpm 45 sec); for PMMA films, 100 µL 5% PMMA in chloroform (Tongguang, 112048, China) containing 2 µM dyes were applied to coverslip followed by spin coating (80,000 rpm 10 sec; 800,000 rpm 45 sec). These polymer films were then irradiated using a Carl ZEISS LSM880 Confocal Microscope equipped with a 20x Air objective (Plan Apochromat 20X/0.8 NA). Three groups of time-lapse images (500 frames,10.6 µm x 10.6 µm, 1.6 fps, 100% laser power) were acquired for each dye, and quantification of the fluorescence intensity was achieved via Analyze >> Tools >> ROI manager in the Fiji software from three parallel experiments.

**Measurement of singlet oxygen quantum yield**
Singlet oxygen quantum yields were determined using 1,3-Diphenylisobenzofuran (DPBF, Q105708, Dibai Shanghai, China) as a chemical indicator. 10 µM dye and 8 x $10^{-5}$ M DPBF were dissolved in acetonitrile (ACN), followed by irradiation with an LED lamp (520-530 nm, 50 mW/cm$^2$). For PKMO, PKMO 0.9, and TMRE, the dyes were corrected for concentration so that their absorbance values at 525 nm were equal. DPBF absorption was monitored at 415 nm, and the linear decay slope of 415 nm is positively correlated with singlet oxygen quantum yield, TMRE ($\varphi$ = 0.012) in MeOH was used as a reference (2).

**Cell culture and isolation of primary cells and tissues**
COS-7 (HARVEYBIO, MB5841) and HeLa (a gift from Prof. Wulan Deng's lab at PKU) cells for phototoxicity measurements were cultured in a high-glucose Dulbecco´s modified Eagle´s medium DMEM medium (Gibco, 11965092) containing 10% (v/v) heat-inactivated fetal bovine serum (VISTECH, SE100-011) and 1% (v/v) penicillin sulfate and streptomycin (Macgene, CC004).
For super-resolution imaging, HeLa cells (3) were grown in DMEM with glutaMAX™ additive and 4.5 g/l glucose (Thermo Fisher Scientific, Waltham, MA, USA). The culture medium was supplemented with 1% (v/v) penicillin-streptomycin (Sigma Aldrich, Munich, Germany), 1 mM sodium pyruvate (Sigma Aldrich), and 10% (v/v) FBS (Merck Millipore, Burlington, MA, USA). U-2 OS cells (European Collection of Authenticated Cell Cultures, ECACC, cat. no. 92022711) were cultured in McCoy´s Medium (Thermo Fisher Scientific) supplemented with 10% (v/v) FBS (Merck Millipore), 1% (v/v) sodium pyruvate (Sigma Aldrich) and 1% penicillin-streptomycin (Sigma Aldrich).
The procedures of primary brown adipocytes, hippocampal neurons, cardiomyocytes, and islet tissues were referenced respectively from (4-7). Primary rat cardiomyocytes (CMs) and primary brown

adipocytes (pBAcs) were cultured in DMEM. Islets tissues isolated from Ins1-Cre$^{+/+}$; GCaMP6f$^{fl/fl}$ mice were cultured on glass-bottom dishes in an RPMI islet medium (RPMI 1640 supplemented with 10% FBS, 8 mM glucose, and 100 U/mL and 100 mg/ml Pen/Strep). Ins1-Cre$^{+/+}$; GCaMP6f$^{fl/fl}$ mice were crossbred by the Ins1-Cre (Jackson Laboratories, stock number 026801) and GCaMP6f$^{fl/fl}$ lines (Jackson Laboratories, stock number 029626) (7). All cells were cultured in an incubator at 37°C with 5% $CO_2$.

**Transfection of cells**

For expression of fusion proteins, HeLa cells were transfected 24-48 h prior to imaging using jetPRIME® transfection reagent (Polyplus-transfection SA, Illkirch-Graffenstaden, France). Transfection was carried out according to the manufacturer protocols using 1-2 µg plasmid DNA. COS-7 cells were transfected 48 h prior to imaging using lipofectamine® 3000 reagents (Thermo Fisher Scientific). Transfection was carried out according to the manufacturer's protocols using 2.5 µg plasmid DNA.

**Plasmids**

CJ-labeling was carried out by transfection with pH-MINOS1-SNAP (8). ER-labeling was performed by transfection with the plasmid Halo-KDEL (9) TOM20-Halo was expressed by transfection with plasmid TOM20-Halo.To generate TOM20-Halo, the plasmid TOM20-eDHFR: L28C (10) was first linearized using the restriction endonucleases EcoRV and XhoI. The gene encoding HaloTag® was amplified by PCR from EF1α-H2B-Halo (a gift from Prof. Wulan Deng's lab) using the primers given below and subsequently integrated into the linearized plasmid by Gibson assembly®.
Halo-fwd: 5'-AGATGATATCAAGCTTACCATGGGCAGCGAAATTGGCACA-3'
Halo-rev: 5'-GCCGCTAATCTCCAGTGTACTCATTACCGCCGCTCCAGAA-3'

**Phototoxicity assay of HeLa cells**

HeLa cells were grown to 80-90 % confluency in a 96-well plate (costar 3599, Corning, NY, USA) before treatment. To achieve the same brightness after staining with two different dyes, HeLa cells were incubated with 1000 nM PKMO and 650 nM PKMO 0.9 respectively in DMEM (Gibco, 11965092) for 10 min at 37℃ with 5% $CO_2$. Then, the cells were washed with PBS three times and maintained in fresh medium for subsequent imaging analysis. 96-well plates were analyzed using a high-content imaging system ImageXpress Micro XLS (Molecular Devices, CA, USA) equipped with a 20 ×/0.4 NA air objective and live-cell imaging device. The average light intensity on cells was measured to be 2.6 W/cm$^2$ (568 nm). The cells in each well were irradiated with the maximum light intensity at different time points. Time points were selected as 5, 7.5, 9, 12.5, 20, and 26.7 min (each time point with three parallel repeats). After illumination, the cells were directly used for subsequent imaging analysis. The cells were washed with PBS once and a 100 µL cell viability assay solution containing 1 µM Calcein AM (Beyotime, C2012-0.1ml, China) was added to each experimental and control group (no dye incubation), then images of two channels (mitochondria: Cy3 channel,568nm, Calcein AM: FITC channel,488nm) were recorded on ImageXpress Micro XLS. The cell viability (%) was calculated according to the following equation: cell viability % = B / A *100%, where A = the number of total cells before illumination, and B = the number of Calcein AM-positive cells after illumination. More than 500 cells were counted at each time point.

**Staining of cell lines, live primary cells, and tissues for STED nanoscopy**

Primary brown adipocytes (pBAcs) were seeded in a 3.5 cm bottom glass dish (STGBD-035-1, Standard Imaging, China) 3 days before imaging measurements. pBAcs and primary hippocampal neurons were stained with DMEM containing 250 nM PKMO at 37 ℃ for 15 min. After removing the staining solution, the cells were then washed with DMEM once and maintained in fresh DMEM for subsequent STED imaging. CMs were seeded in laminin-coated glass-bottom dishes 3 days before measurements. CMs were stained with 500 nM PKMO in DMEM for 15 min before imaging experiments. Islets tissues isolated from Ins1-Cre+/+; GCaMP6ffl/fl mice (7) were stained with Krebs-Ringer bicarbonate buffer (KRBB) solution containing 125 mM NaCl, 5.9 mM KCl, 2.4mM $CaCl_2$, 1.2mM $MgCl_2$, 1 mM L-Glutamine, 25 mM HEPES, 3 mM glucose, 0.1% (v/v) bovine serum albumin, and 600 nM PKMO at 37 ℃ for 30 min. After removing the staining solution, the islets were then washed with KRBB solution three times and maintained in a fresh medium for the STED imaging.

COS-7 cells were seeded in glass-bottom dishes (STGBD-035-1, Standard Imaging, China) two days prior to imaging. COS-7 cells were stained with DMEM supplemented with 250 nM PKMO for 15 min. U-2 OS cells and HeLa cells were seeded in glass-bottom dishes (ibidi GmbH, Germany) one day prior to imaging. U-2 OS cells were stained with McCoy´s medium supplemented with 250 nM PKMO for 20 min. For single-color recordings, HeLa cells were stained with DMEM containing 150-250 nM PKMO for 40-45 min. Following the staining procedure, cells were washed three times with culture medium and incubated for 30-60 min to remove the unbound dye. The cells were imaged at room temperature in HEPES-buffered DMEM (HDMEM) containing 4.5 g/l glucose, l-glutamine, and 25mM HEPES (Thermo Fisher Scientific).

**Sample preparation for multi-color imaging**

*Labeling of cristae and mtDNA in HeLa cells*
For labeling of cristae and mtDNA, HeLa cells were incubated with DMEM containing 150-300 nM PKMO for 45-60 min. Cells were washed twice and incubated with DMEM containing 0.5 µl/ml Quant-iT PicoGreen reagent (Thermo Fisher Scientific) for 20 min. Afterward, the culture medium was replaced and the cells were incubated for 30 min to remove unbound dye. Cells were recorded in HDMEM at room temperature.

*Labeling of cristae and SNAP/Halo fusion proteins*
For labeling of self-labeling tags and cristae, HeLa cells were incubated with DMEM containing 200-300 nM PKMO and 0.5 µM 647-SiR-CA (for Halo-KDEL) or 1 µM SNAP-Cell 647-SiR (for MIC10-SNAP; New England BioLabs Inc., Ipswich, MA, USA) for 40-45 min. Following the staining procedure, cells were washed three times with culture medium and incubated for approximately 60 min to remove the unbound dye. The cells were imaged in HDMEM at room temperature. COS-7 cells were incubated with DMEM containing 500 nM 647-SiR-CA (for TOM20-Halo, and synthesized according to literature protocol (11) ) for 10 min. After removing the staining solution and three washing steps with PBS, the cells were then stained with DMEM containing 250 nM PKMO for 15 min at 37 ℃. After removing the staining solution, the cells were then washed with DMEM once and maintained in fresh DMEM for following STED imaging.

*Labeling of cristae, tubulin, and mtDNA.*
For labeling of cristae together with cytoskeleton and mtDNA, HeLa cells were incubated with DMEM containing, 350 nM PKMO, 200 nM 4-610CP-CTX (12), and 0.2 µl/ml Quant-iT PicoGreen reagent (Thermo Fisher Scientific) for 40 min. Cells were washed three times with culture medium and incubated for 60 min to remove unbound dye. Cells were recorded in HDMEM at room temperature.

**Live-cell imaging of cancer cells, primary cells, and tissues**
COS-7 cells, primary cells, and tissues were recorded using an Abberior Facility Line and Abberior STEDYCON (Abberior Instruments GmbH, Göttingen, Germany) fluorescence microscope equipped with an Olympus UPlanXAPO 60x, NA1.42 objective (Olympus, Tokyo, Japan). Pixel sizes of 20-30 nm were used for STED nanoscopy. PKMO was excited at 561 nm wavelength and STED was performed using a pulsed depletion laser at 775 nm wavelength, with gating of 1-7 ns and dwell times of 10 µs. For STED images, each line was scanned 3 to 5 times and the signal was accumulated. For the confocal images, each line was scanned once. For one-color STED imaging of live COS-7 cells, all images were recorded using Facility Line except for the data in figure 2d and figure S1b-c were recorded using STEDYCON. For dual-color STED imaging of COS-7 cells, 647-SiR-CA was excited at 640 nm wavelength. STED was performed at 775 nm wavelength with gating of 0.75-8.75 ns. Dwell times of 10 µs were used. Each line was scanned 3 times and the signal was accumulated. 3D STED nanoscopy of COS-7 cells was carried out using a 3D STED PSF of Facility Line. The voxel size was 20 x 20 x 50 nm for one-color 3D STED and 20 x 20 x 30 nm for dual-color 3D STED.

STED nanoscopy of HeLa and U-2 OS cells was carried out using an Expert Line dual-color STED 775 QUAD scanning microscope (Abberior Instruments GmbH). The microscope was equipped with a UPlanSApo 100×/1.40 Oil [infinity]/0.17/FN26.5 objective (Olympus). In brief, PKMO was excited at 561 nm and SiR was excited at 640 nm wavelength. Depletion was performed at a 775 nm wavelength. PicoGreen was excited at 488 nm wavelength and was recorded in the confocal mode. Imaging parameters were individually adapted for each sample. Typically, pixel sizes of 20 - 30 nm and dwell times of 5-7 µs were used. In the STED mode, each line was scanned 6 to 9 times and the signal was accumulated. The pinhole was set to 0.7 to 1.0 AU. The signal gating was set to 0.5 – 8 ns.

**Electron microscopy of HeLa cells**

Sample preparation was performed according to Stephan et al., 2020 (8). In brief, HeLa cells were grown on Aclar® discs to a confluency of approximately 70%. Fixation was performed by immersion with pre-warmed (37 °C) 2% glutaraldehyde in 0.1 M cacodylate buffer (pH 7.4) at room temperature. For complete fixation, samples were stored at 4°C overnight. Following post-fixation with 1% osmium tetroxide and pre-embedding staining with 1% uranyl acetate, samples were dehydrated and resin-embedded. Ultrathin sections were recorded on a Talos L120C transmission microscope (Thermo Fischer Scientific, Hilsboro, Oregon, USA) at 11,000-13,500× magnification using a Ceta 4k × 4k CMOS camera.

**Image processing**

*STED microscopy*

STED nanoscopy images of COS-7 cells, primary cells, and tissues were deconvoluted using Huygens software (Scientific Volume Imaging B.V., Hilversum, The Netherlands). Signal to Noise Ratio (SNR) was estimated by Huygens software. STED nanoscopy images of HeLa and U-2 OS cells and 3D STED images of COS 7 cells were deconvoluted using the Richardson-Lucy algorithm in the Imspector software (Abberior Instruments GmbH; version 0.14.11616). 3D rendering and reconstruction of 3D STED data were performed using Imaris (Bitplane, Belfast, UK).

Resolution estimation was performed by the Gaussian fitting of fluorescence intensity line profiles. The full width at half maxima (FWHM) was estimated via Analysis >> Fitting >> Nonlinear curve fit >> Gaussian fit in the Origin Pro 2020b software (OriginLab Corporation, Northampton MA, USA) or using Matlab (The Math Works, Natick, MA, USA). All images that were used to analyze the resolution in supplementary information are raw data without background subtraction. For time-lapse recordings, photobleaching was compensated using the bleach correction feature in Fiji/ImageJ (version 1.53f51).

*Electron microscopy*

Electron microscopy recordings were filtered using a median filter in Fiji/ImageJ (version 1.53f51).

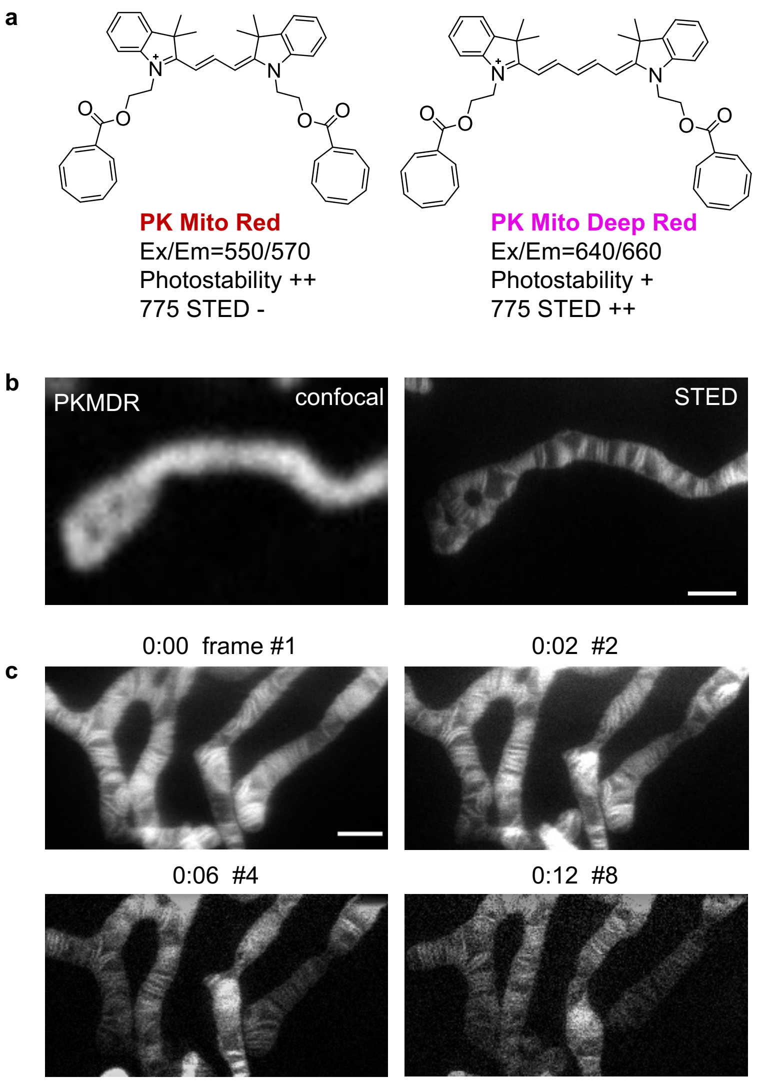


**Figure S1. STED nanoscopy of mitochondria labeled with PKMDR.** (a) Chemical structures of PK Mito Red (PKMR) and PK Mito Deep Red (PKMDR). (b) Confocal (left) and STED (right) images of mitochondria in live COS-7 cells labeled with 250 nM PKMDR for 15 min. Scale bar = 1 µm. (c) Time-lapse images of cristae in a live COS-7 cell labeled with PKMDR (250 nM, 15 min). Scale bar = 1 µm.

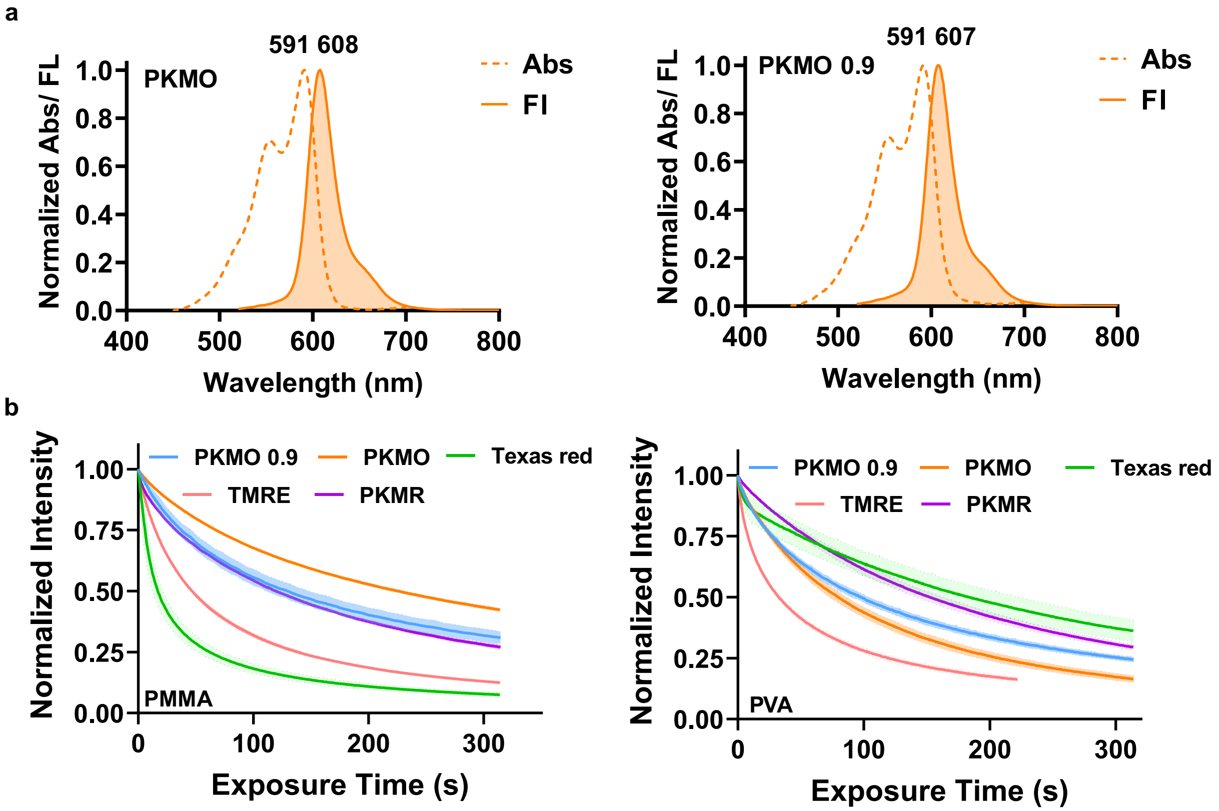


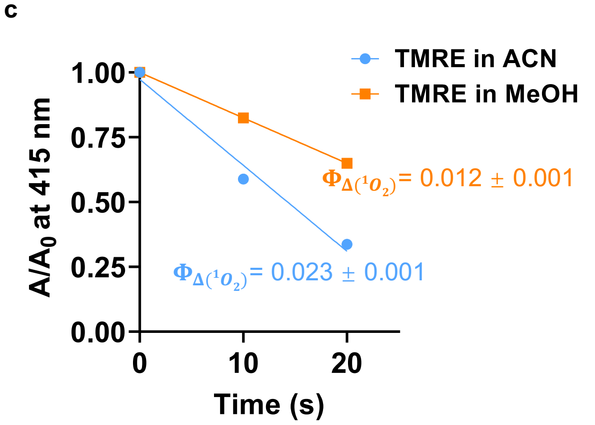


d

| | PKMO | PKMO 0.9 |
|---|---|---|
| **Abs/Em** | 591/608[a] | 591/607[a] |
| **Quantum yield ($\Phi_f$)** | 0.14[a] / 0.45[b] | 0.13[a] / 0.49[b] |
| **$\Phi_{\Delta(^1O_2)}$ (x $10^{-4}$)** | 67 ± 5[c] | 17 ± 1[c] |

**Figure S2. Photophysical properties of PKMO.** (a) Absorption (abs) and fluorescence (fl) spectra of PKMO (left) and PKMO 0.9 (right) in methanol. (b) Photobleaching curve of PKMO, PKMO 0.9, PKMR, Texas Red, TMRE in polymethyl methacrylate (PMMA) (left) and polyvinyl alcohol (PVA) film (right) recorded under continuous 561 nm laser scanning of a confocal microscope. (c) Singlet oxygen quantum yield of TMRE in MeOH or ACN measured using 1,3-diphenylisobenzofuran (DPBF) decay assay. TMRE in MeOH ($\Phi_\Delta$= 0.012) in MeOH was selected as a standard. (d) table of photophysical properties of PKMO and PKMO 0.9. Solvents: a, methanol; b, toluene; c, acetonitrile.

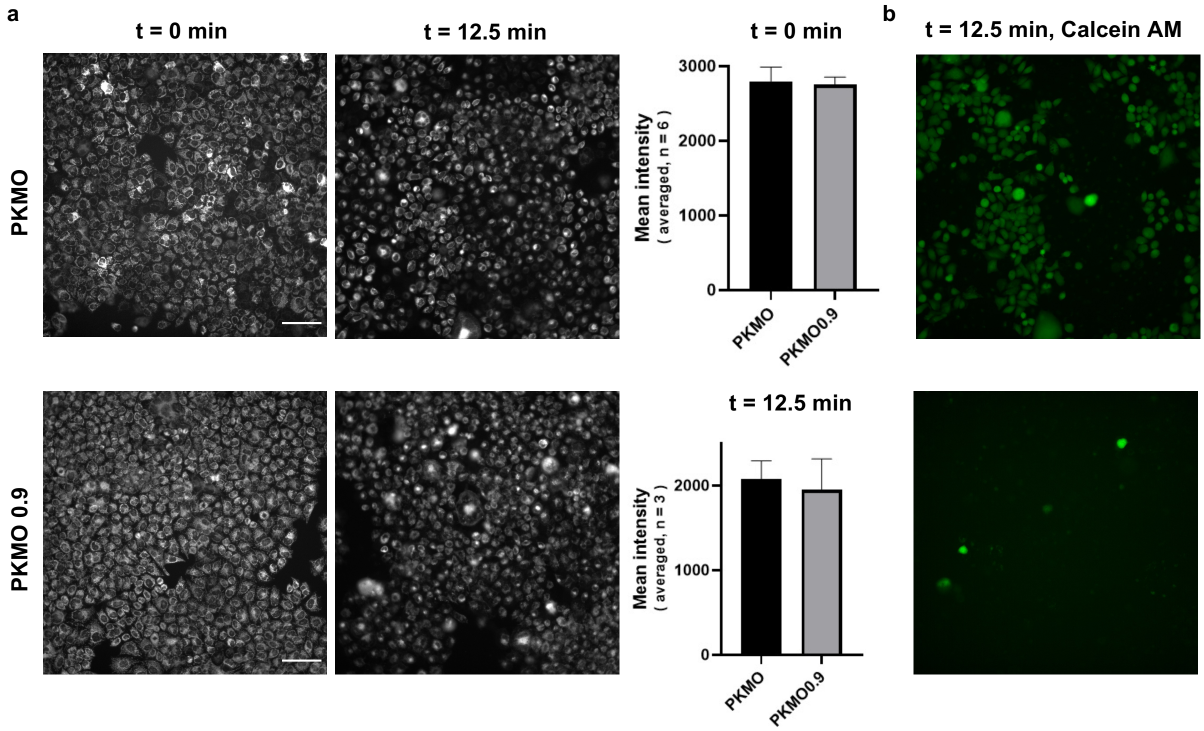


**Figure S3. Phototoxicity of PKMO and PKMO 0.9.** (a) Wide field images of live HeLa cells labeled with 1 μM PKMO or 650 nM PKMO 0.9 for 15 min before (left) and after LED illumination in the high-content imaging system and mean intensity analysis of mitochondria before and after 12.5-min illumination. (b) Wide field images of Calcein AM (2 μM) indicating live cells labeled with PKMO and PKMO 0.9 after 12.5-min illumination.

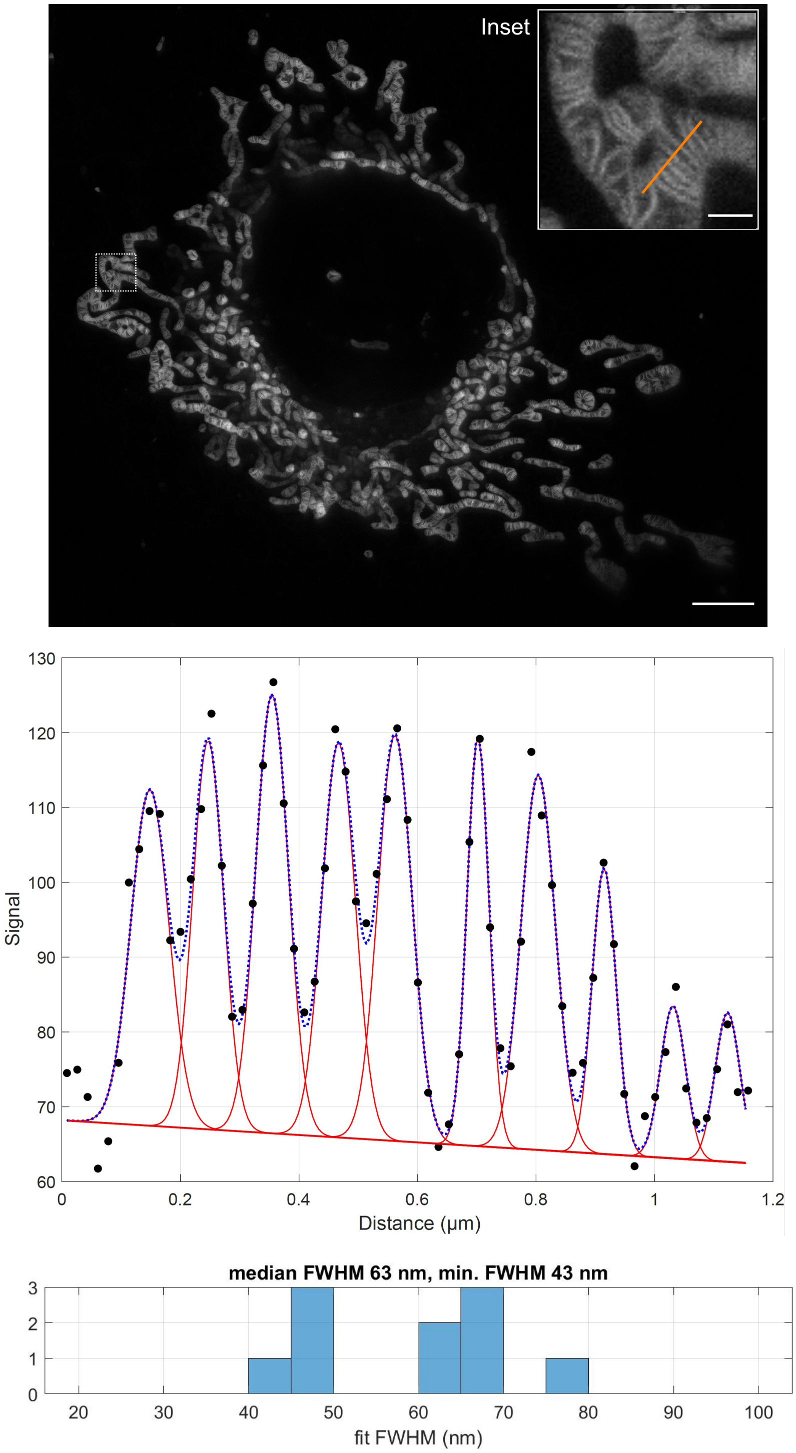


**Figure S4. Resolution estimation of 2D STED nanoscopy.** Top: raw data (SNR=15) of the 2D-STED recording of a live COS-7 cell shown in Fig. 2. Inset (white boxed area) indicates the area used for resolution estimation. Bottom: fluorescence intensity line profile was recorded along the orange line shown in the inset. The fluorescence intensity signal was fitted using a Gaussian fit. The full width at half maximum (FWHM) was estimated for the individual peaks. Scale bar: overview 5 µm, inset 2 µm.

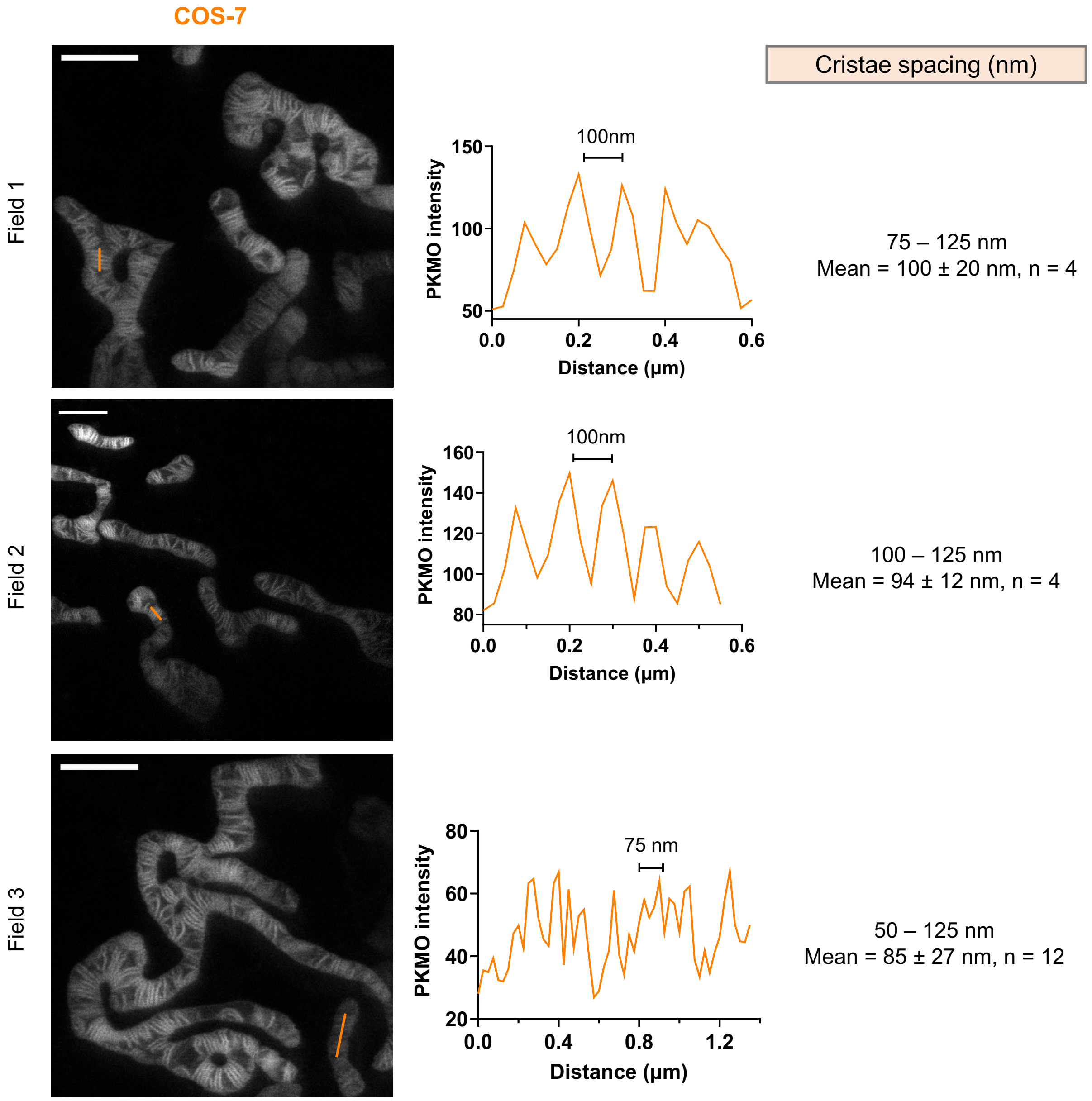


**Figure S5. Cristae spacing in COS-7 cells.** STED images of mitochondrial cristae in live COS-7 cells labeled with PKMO (left) and fluorescence intensity line profiles (right). The fluorescence signal was measured as indicated by the orange lines in the STED images and cristae distances were estimated. Scale bars = 2 µm.

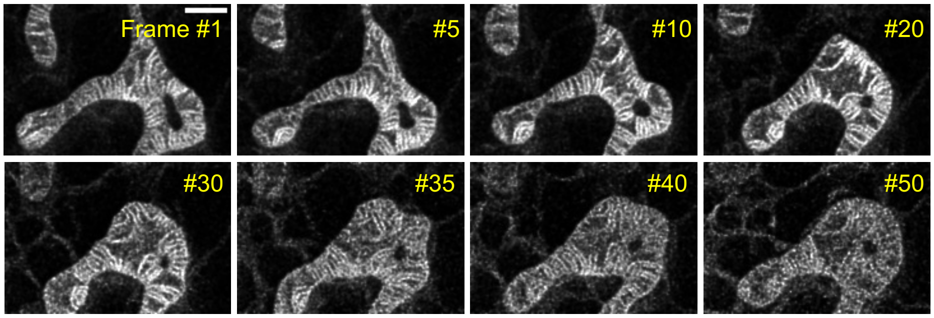


**Figure S6. Time-lapse 2D STED nanoscopy of COS-7 cells.** Shown are selected frames from a 50-frame STED recording of cristae in a live COS-7 cell labeled with PKMO (250 nM,15 min). Scale bar = 1 μm.

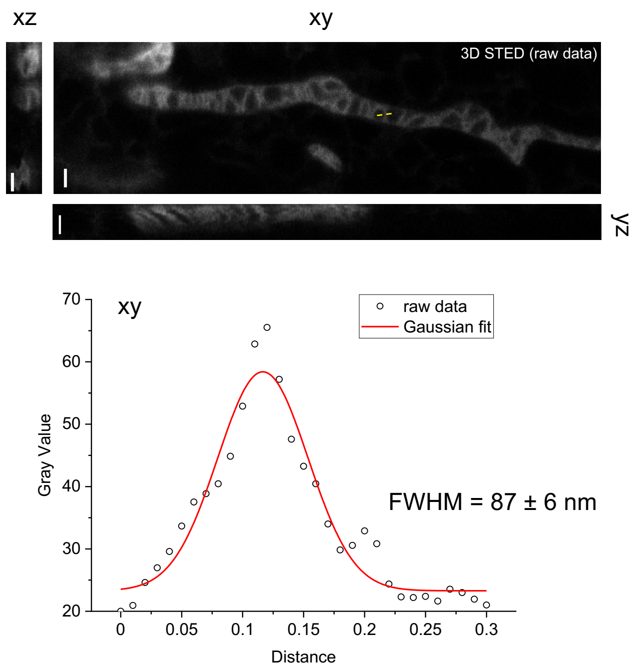


**Figure S7. 3D STED nanoscopy of COS-7 cells.** Orthogonal view of a 3D-STED recording of a mitochondrion in a live COS-7 cell presented in Fig. 3. The image was recorded using a 3D STED PSF. Voxel size 20 x 20 x 50 nm. The fluorescence intensity line profile was measured as indicated by the yellow line. Fluorescence intensity was fitted using a Gaussian fit and FWHM was estimated. Scale bars = 500 nm.

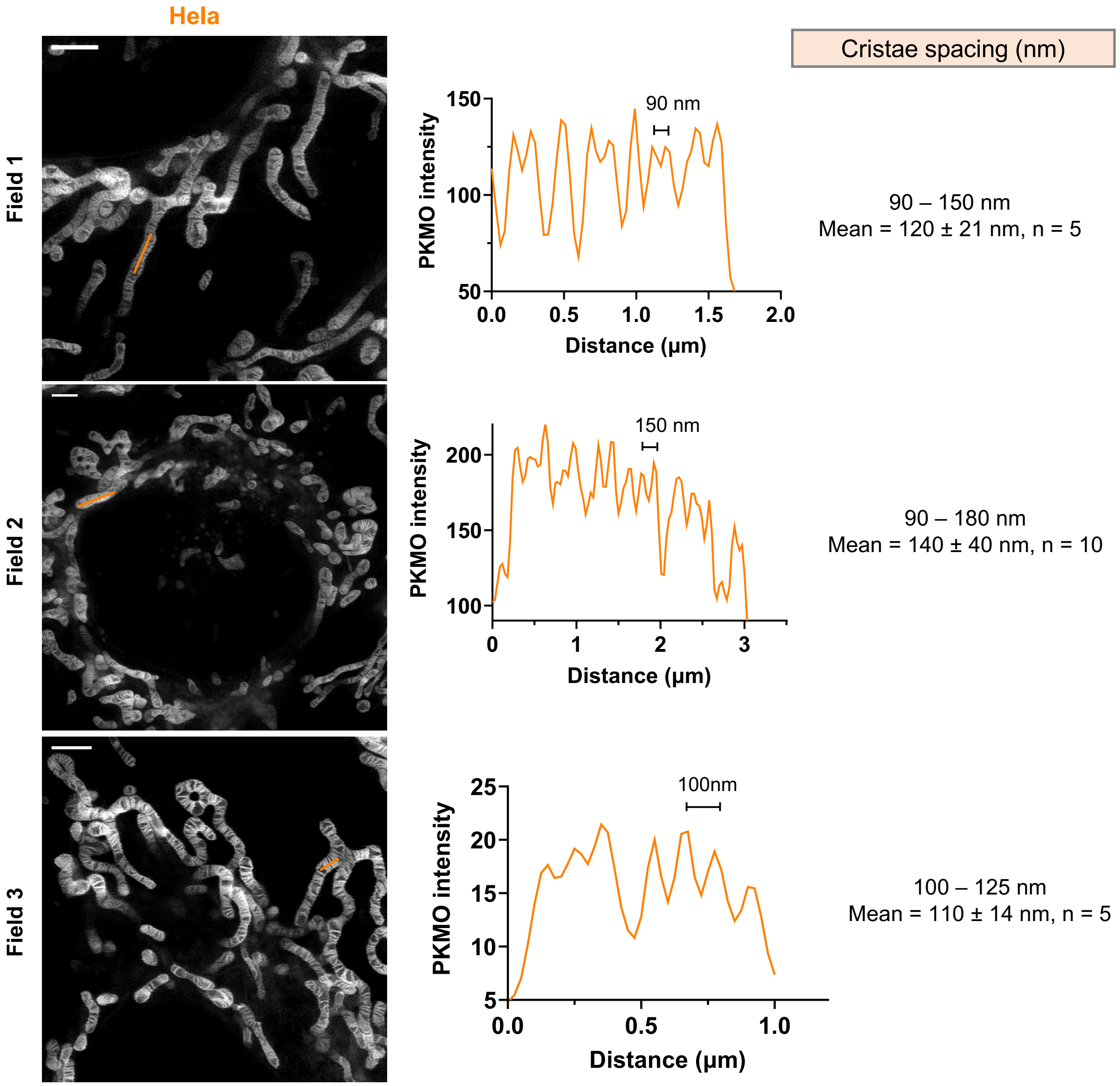


**Figure S8. Cristae spacing in HeLa cells.** STED images of mitochondrial cristae in live HeLa cells labeled with PKMO (left) and fluorescence intensity line profiles (right). The fluorescence signal was measured as indicated by the orange lines in the STED images and cristae distances were estimated. Scale bars = 2 µm.

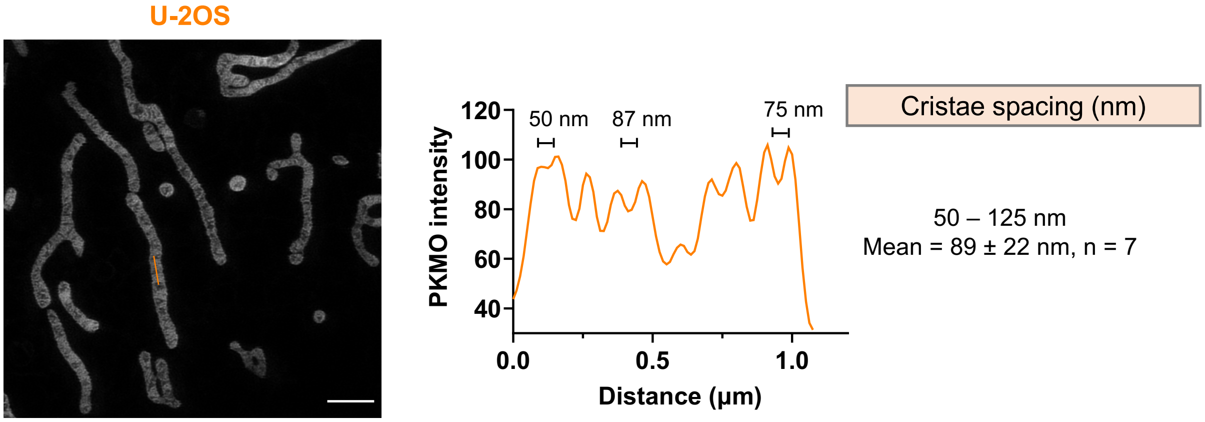


**Figure S9. Cristae spacing in U-2 OS cells.**
STED image of mitochondrial cristae in a live U-2 OS cell labeled with PKMO (left) and fluorescence intensity line profiles (right). The fluorescence signal was measured as indicated by the orange line in the STED image and cristae distances were estimated. Scale bars = 2 µm.

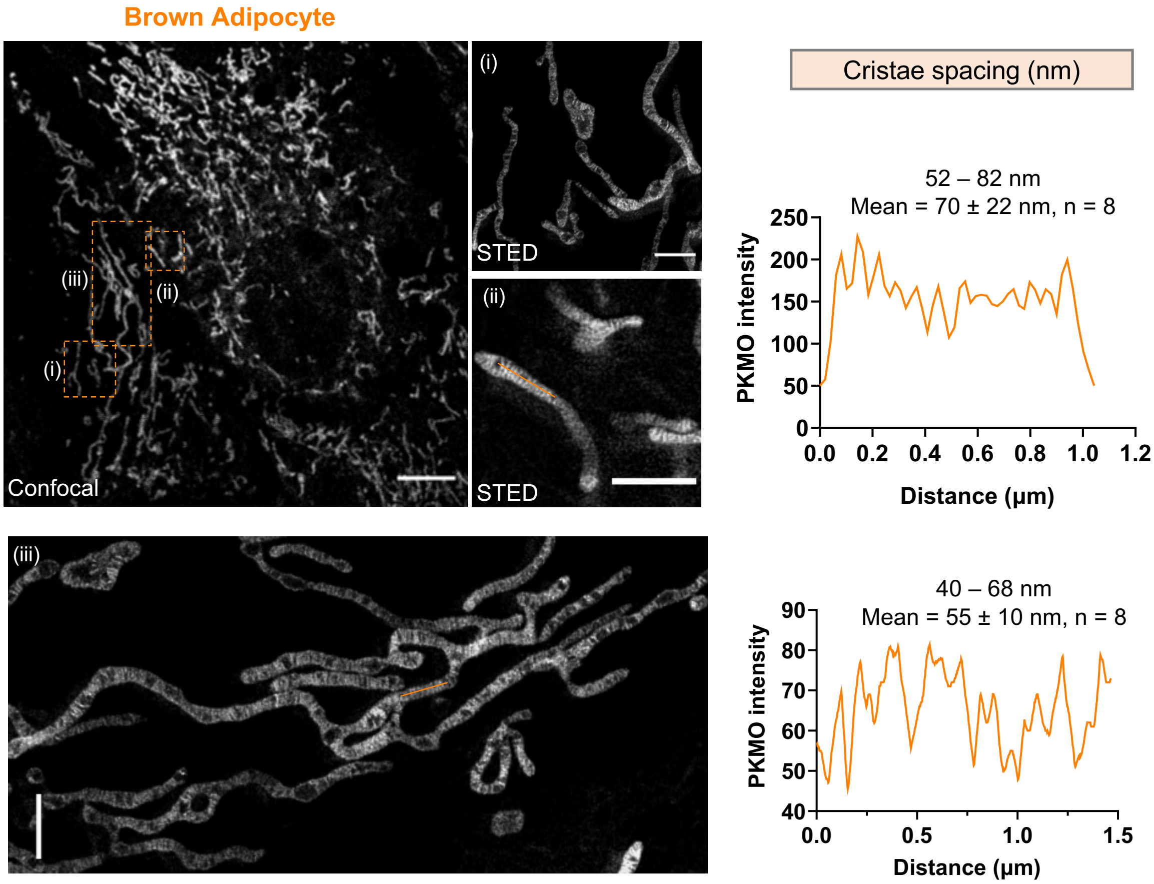


**Figure S10. Cristae spacing in brown adipocytes.** STED images of mitochondrial cristae in live primary brown adipocytes (pBAcs) labeled with PKMO (250 nM,15 min) (left) and fluorescence intensity line profiles (right). The fluorescence signal was measured as indicated by the orange lines in the STED images and cristae distances were estimated. Scale bar: overview 5 µm, insets 2 µm.

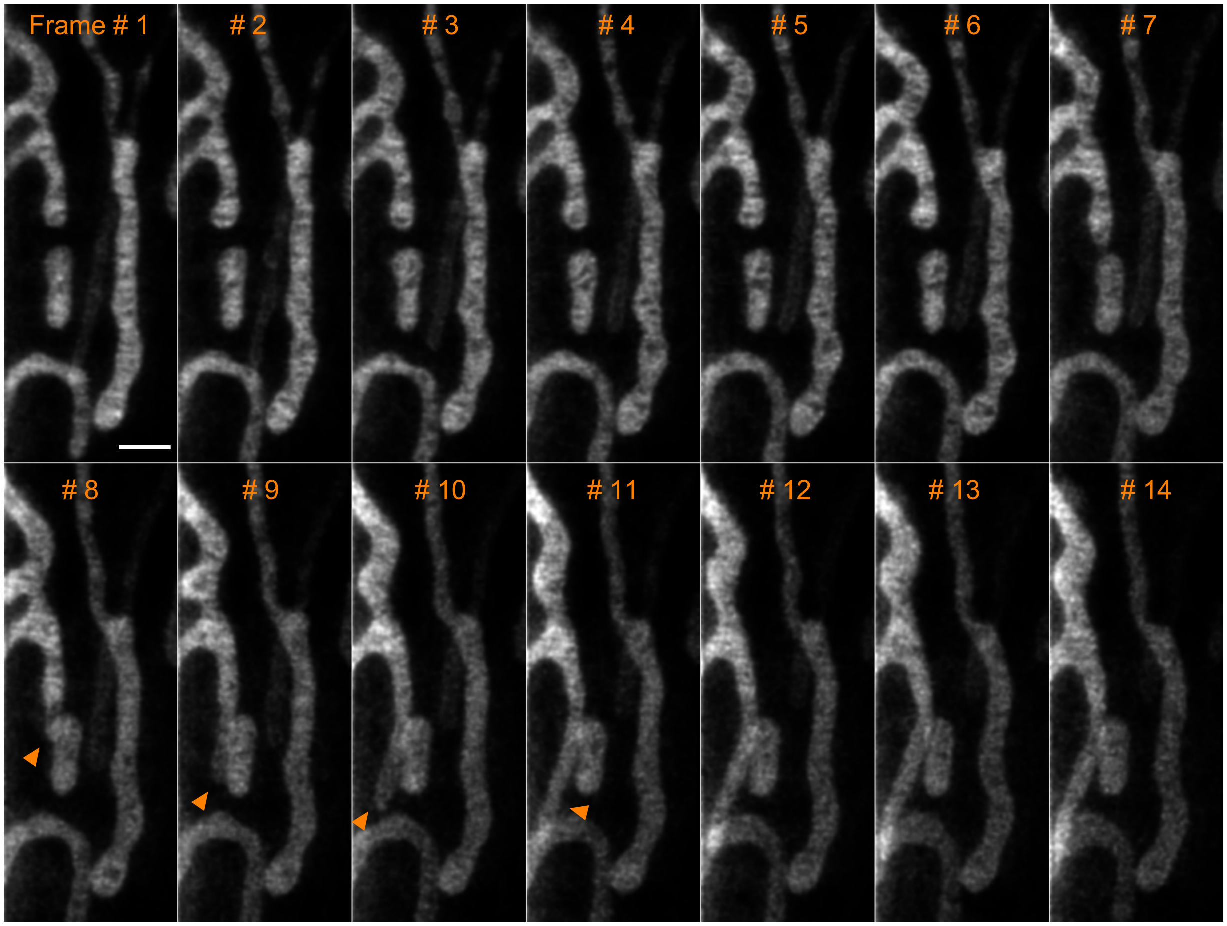


**Figure S11. Time-lapse STED nanoscopy of brown adipocytes.** Time-lapse STED recording of primary brown adipocytes labeled with PKMO (250 nM staining for 15 min). Orange triangles indicate an event of mitochondrial tubulation. Scale bar = 1 μm.

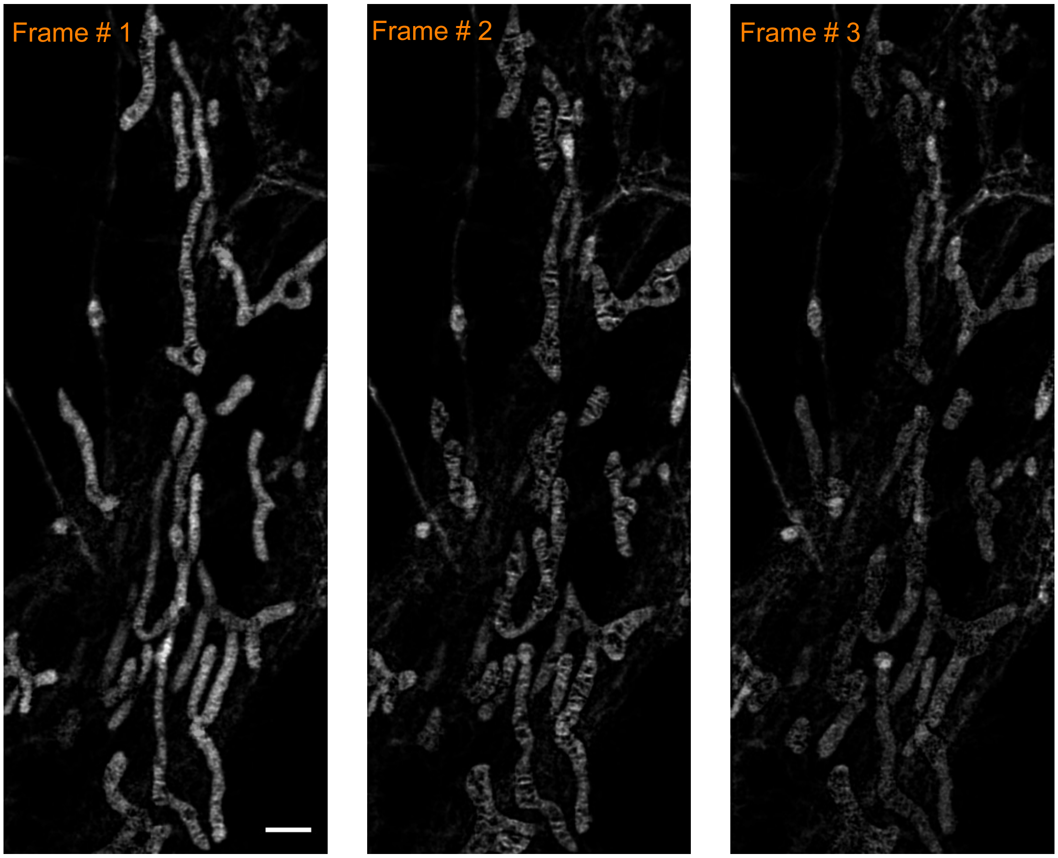

**Figure S12. Time-lapse STED nanoscopy of neurons.** Time-lapse STED recordings of primary mouse hippocampal neurons labeled with PKMO (250 nM, 15 min). Images show photobleaching and structural changes caused by phototoxic effects. Scale bar = 1 μm.

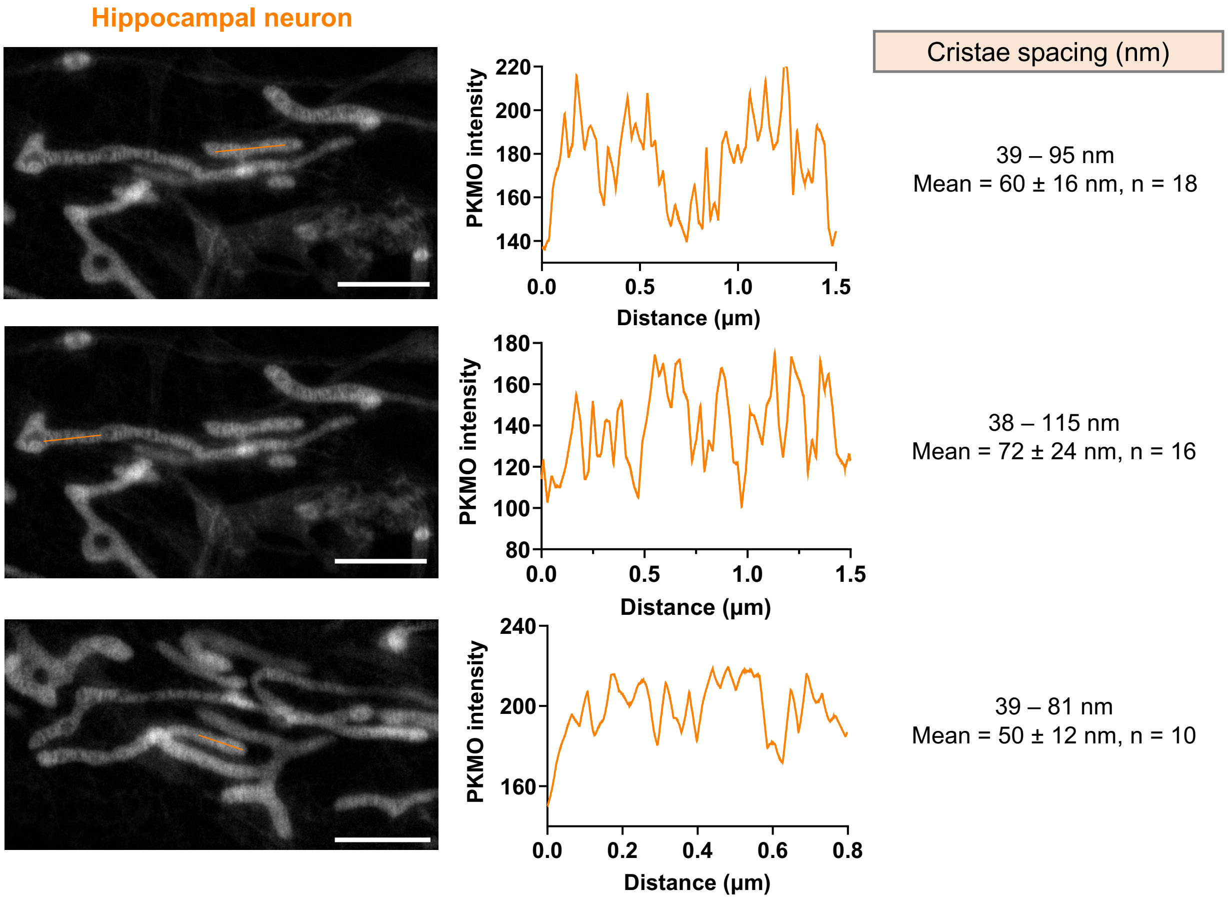


**Figure S13. Cristae distance in neurons.** STED images of mitochondrial cristae in live primary hippocampal neurons labeled with PKMO (left) and fluorescence intensity line profiles (right). The fluorescence signal was measured as indicated by the orange lines in the STED images and cristae distances were estimated. Scale bars = 2 μm.

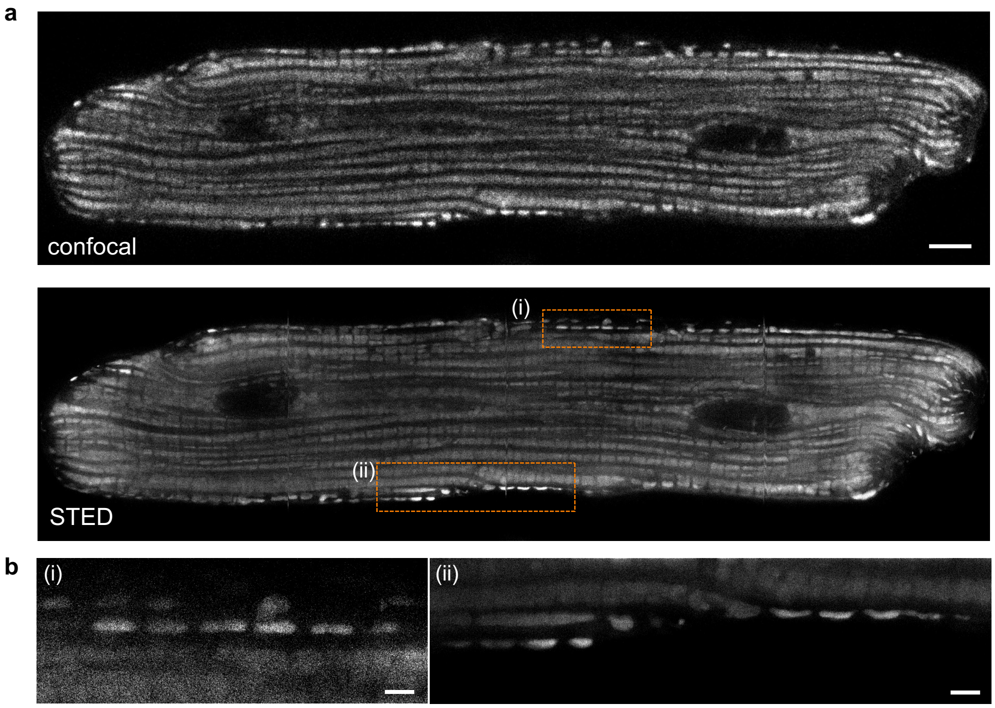


**Figure S14. PKMO labeling in rat cardiomyocytes.** (a) Confocal and STED images of mitochondria in live primary rat cardiomyocytes labeled with PKMO (500 nM, 15 min). Scale bar = 5 μm. (b) Magnifications of the area indicated in the STED overview images in (a). Scale bars = 1 μm.

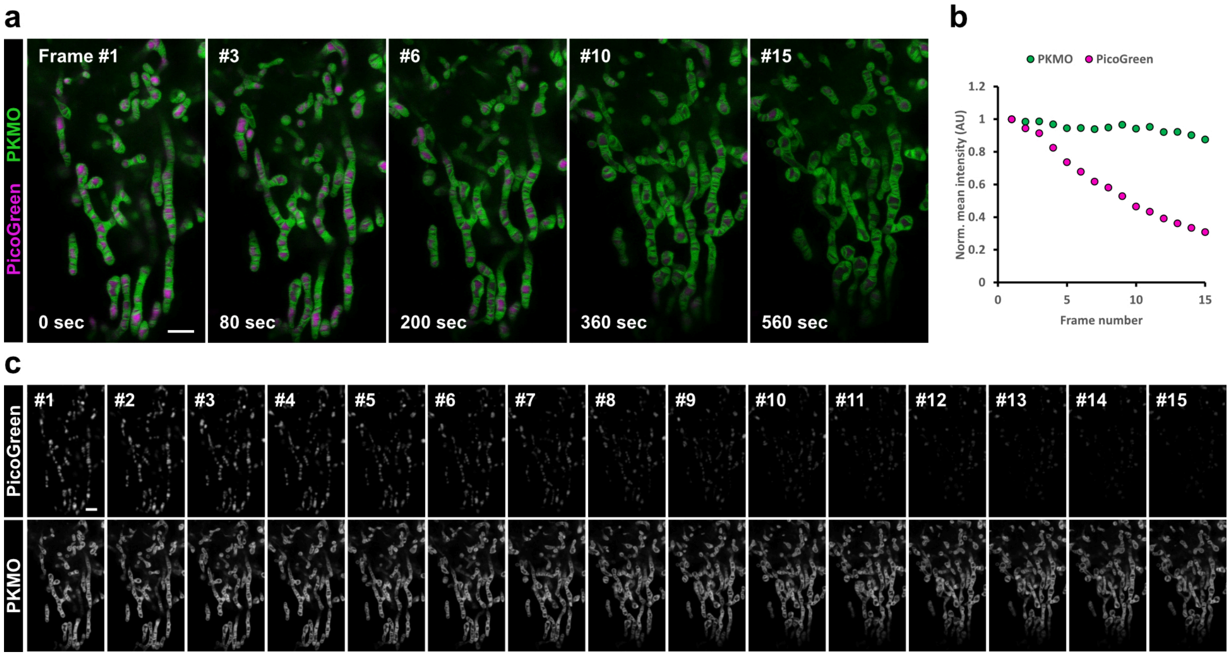


**Figure S15. Photobleaching of PKMO and PicoGreen.** (a-c) 2D dual-color time-lapse recording of mitochondrial cristae (STED) and DNA (confocal). (a) Selected frames of the time-lapse recording presented in Figure 5b. Images illustrate bleaching and changes of mitochondrial morphology over time. (b) Photobleaching during dual-color time-lapse images based on the frames presented in (c). Scale bars = 2 µm.

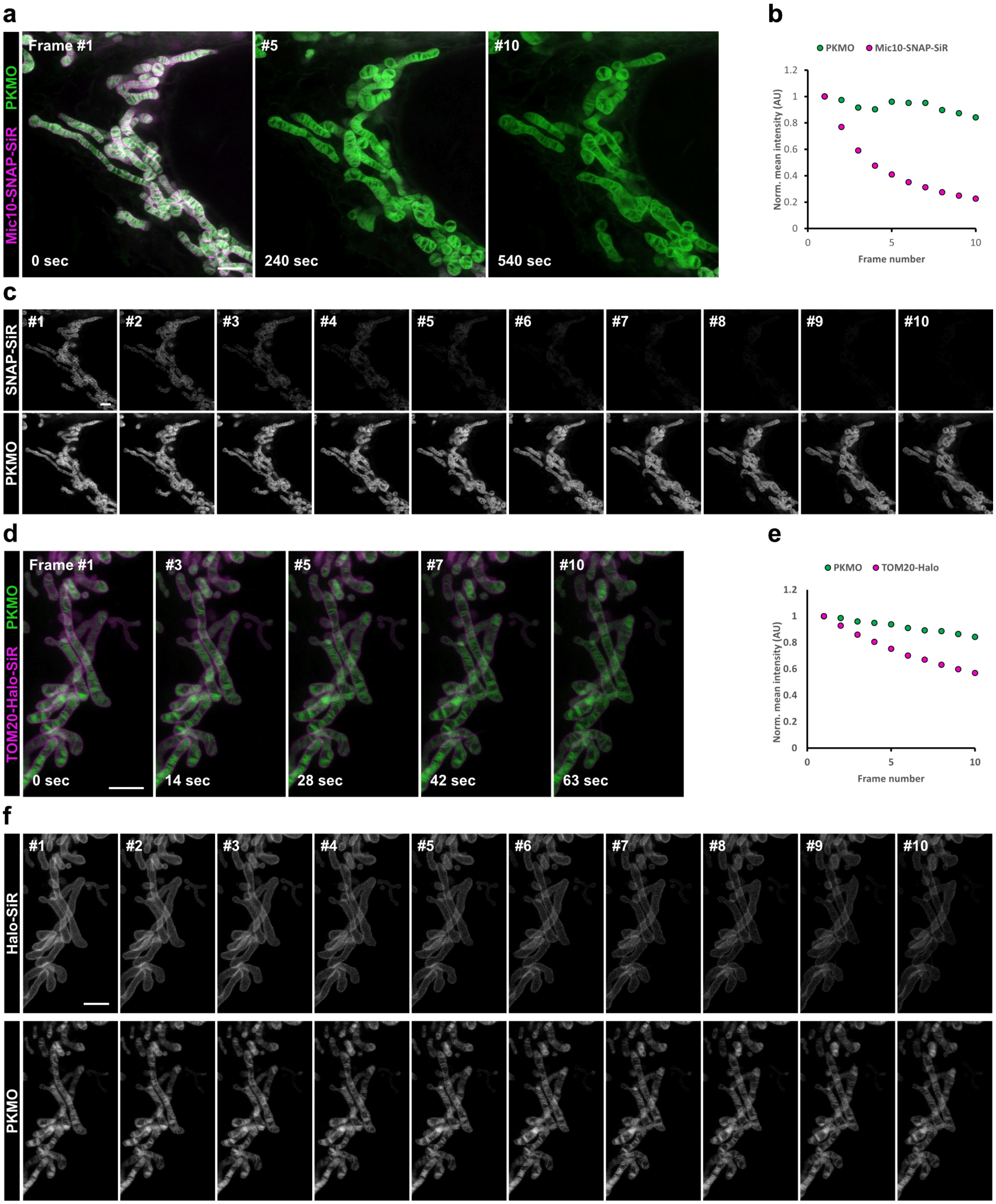


**Figure S16. Photobleaching of PKMO and Mic60-SNAP-SiR or TOM20-Halo-SiR.** (a-c) 2D dual-color time-lapse STED recording of mitochondrial cristae (PKMO) and Mic10-SNAP (SNAP-Cell-647-SiR). (a) Selected frames of the time-lapse recording presented in Figure 5e. Images illustrate bleaching and changes of mitochondrial morphology over time. (b) Photobleaching during dual-color time-lapse imaging based on the frames presented in (c). (d-f) 2D dual-color time-lapse STED images of mitochondrial cristae (PKMO) and TOM20-Halo (647-SiR-CA). (d) Selected frames of the time-lapse recording presented in Figure 5c. Images illustrate bleaching and changes of mitochondrial morphology over time. (e) Photobleaching during dual-color time-lapse imaging based on the frames presented in (f). Scale bars = 2 μm.

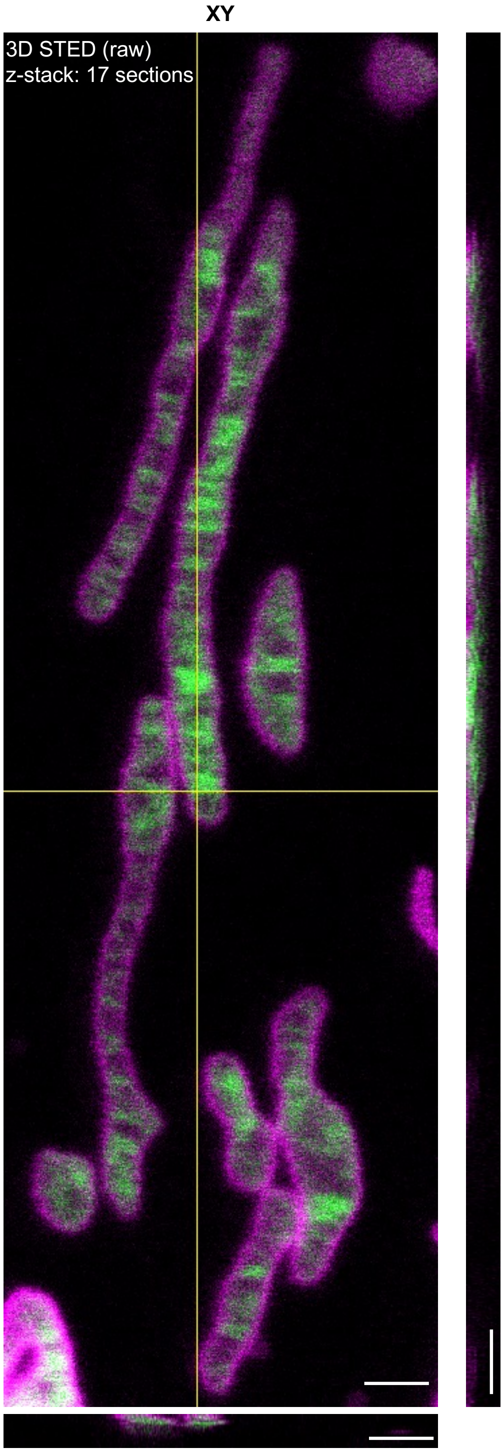


**Figure S17. Dual-color 3D STED nanoscopy of mitochondria.** Orthogonal view of the dual-color 3D-STED recording of mitochondrial cristae (PKMO, green) and outer membrane (TOM20-Halo, magenta) presented in Fig. 5d. Voxel size= 20 x 20 x 30 nm (XYZ). Scale bars = 1 μm.

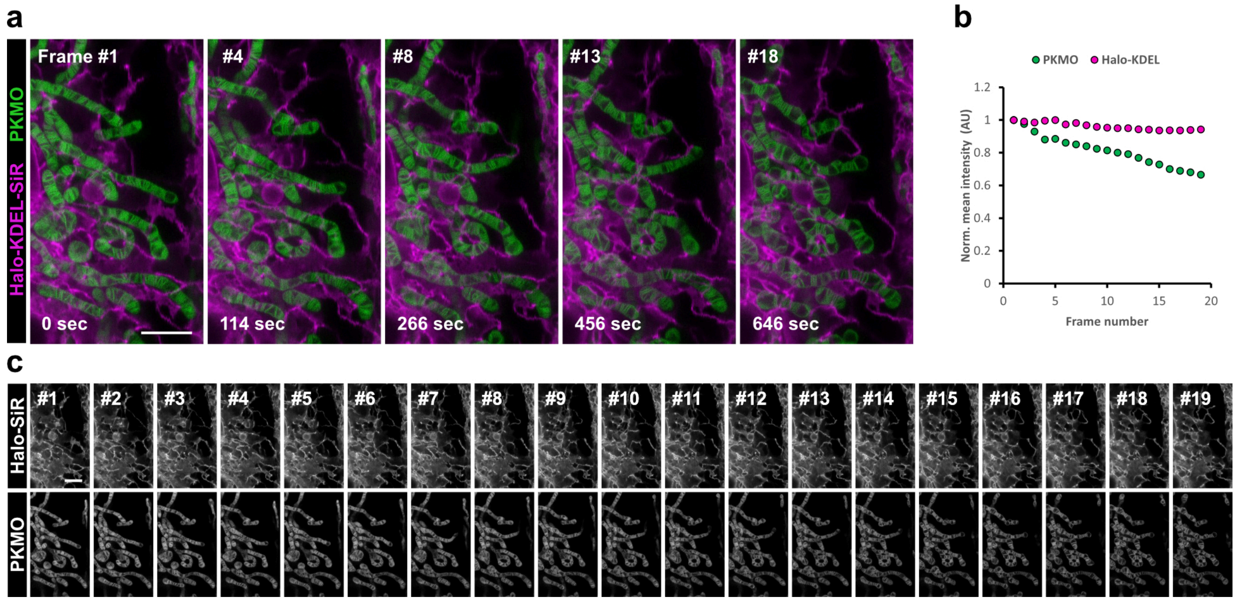


**Figure S18. Photobleaching of PKMO and Halo-KDEL-SiR in HeLa cells.** (a-c) 2D dual-color time-lapse STED recording of mitochondrial cristae (PKMO staining) and Halo-KDEL (647-SiR-CA staining). (a) Selected frames of the time-lapse recording presented in Figure 5e. Images illustrate bleaching and changes of mitochondrial morphology over time. (b) Photobleaching during dual-color time-lapse imaging based on the frames presented in (c). Scale bars = 2 μm.

**Table S1. Comparison of cristae spacing of various cell lines analyzed from STED images.**

| Cells | Cristae spacing range (nm) | Mean ± SD of cristae spacing (nm) |
|---|---|---|
| HeLa | 90 – 150 | 127 ± 33, n = 20 |
| COS-7 | 50 – 125 | 90 ± 24, n = 20 |
| U-2OS | 50 - 125 | 89 ± 22, n = 7 |
| Primary brown adipocytes | 40 - 82 | 63 ± 14, n = 16 |
| Primary hippocampal neurons | 38 - 115 | 62 ± 20, n = 44 |

### Chemical Synthesis and Characterization of New Compounds

#### General Information

Unless otherwise mentioned, all reactions were carried out under a nitrogen atmosphere with dry solvents under anhydrous conditions. All the chemicals were purchased at the highest commercial quality and used without further purification unless otherwise stated. Reactions were monitored by Thin Layer Chromatography on plates (GF254) supplied by Yantai Chemicals (China) using UV light as a visualizing agent and an ethanolic solution of phosphomolybdic acid and cerium sulfate, and heat as developing agents or by LC/MS (4.6 mm × 150 mm 5 μm C18 column; 5 μL injection; 10-95% or 50-95% $CH_3CN/H_2O$, linear-gradient, with constant 0.1% v/v TFA additive; 20 min run; 1 mL/min flow; ESI; positive ion mode; UV detection at 254 nm). Flash column chromatography uses silica gel (200-300 mesh) supplied by Tsingtao Haiyang Chemicals (China). NMR spectra were recorded on Brüker Advance 400 ($^{1}$H 400 MHz, $^{13}$C 101 MHz) and are calibrated using residual undeuterated solvent ($CDCl_3$ at 7.26 ppm $^{1}$H NMR, 77.16 ppm $^{13}$C NMR; $CD_3OD$ at 3.31 ppm $^{1}$H NMR, 49.00 ppm $^{13}$C NMR; DMSO-d6 at 2.50 ppm $^{1}$H NMR, 39.52 ppm $^{13}$C NMR). Data for $^{1}$H NMR spectra are reported as follows: chemical shift (δ ppm), multiplicity (s = singlet, d = doublet, t = triplet, q = quartet, dd = doublet of doublets, dt = triplet of doublets, m = multiplet, br = broad), coupling constant (Hz), integration. Data for $^{13}$C NMR are reported by chemical shift (δ ppm).

#### Synthetic Procedure and Characterization

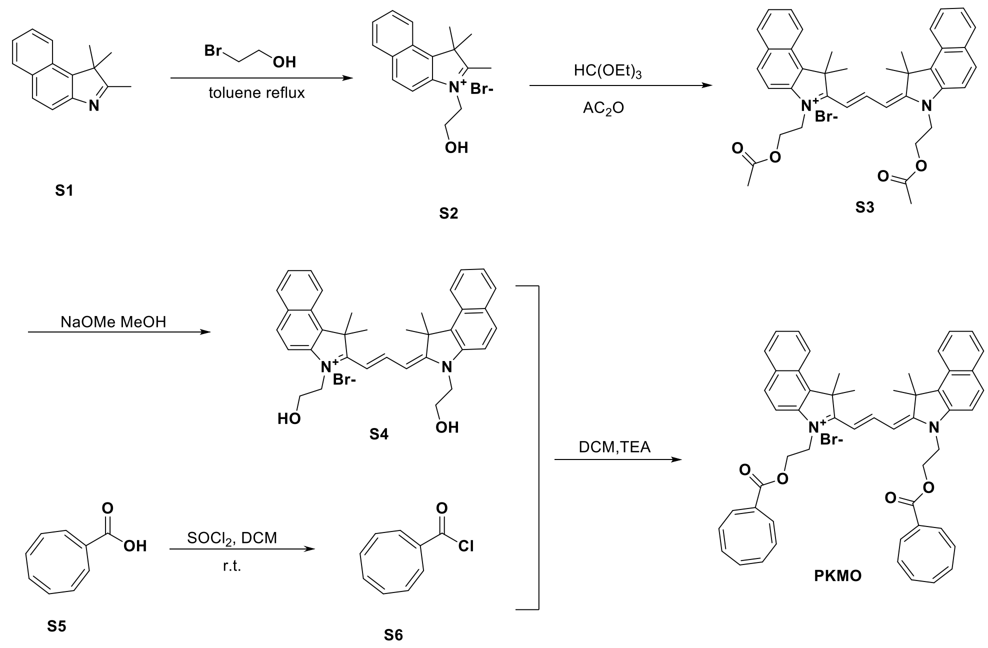


**Scheme S1**. Synthetic route of Compound PKMO and PKMO 0.9

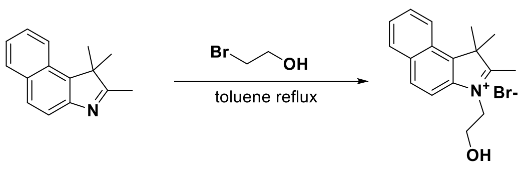


**S1** **S2**

**Compound S2**: To a solution of compound S1 (500 mg, 2.39 mmol) in toluene (5 mL) was added **2-Bromoethan-1-ol** (CAS: 540-51-2) (896 mg, 7.17 mmol) at room temperature (r.t.), the mixture was heated at 110°C and stirred for 16 h. The reaction mixture was then cooled to r.t slowly and a solid precipitated. The solid was collected by filtration, followed by washing with MTBE (10 mL), the solid was then dried under vacuum to afford the compound (700 mg, 88%) as a brown solid.

$^1$H NMR (400 MHz, Methanol-d4) δ 8.34 (d, J = 8.5 Hz, 1H), 8.23 (d, J = 9.0 Hz, 1H), 8.17 (d, J = 8.3 Hz, 1H), 8.01 (d, J = 9.0 Hz, 1H), 7.81 (td, J = 8.5, 1.4 Hz, 1H), 7.72 (td, J = 8.3, 1.2 Hz, 1H), 4.79(t, J =6.0 Hz, 2H), 4.12 (t, J =6.0 Hz, 2H), 1.86 (s, 6H).

MS (ESI) calcd for $C_{17}H_{20}NO^+$ $[M]^+$ 254.15, found 254.06.

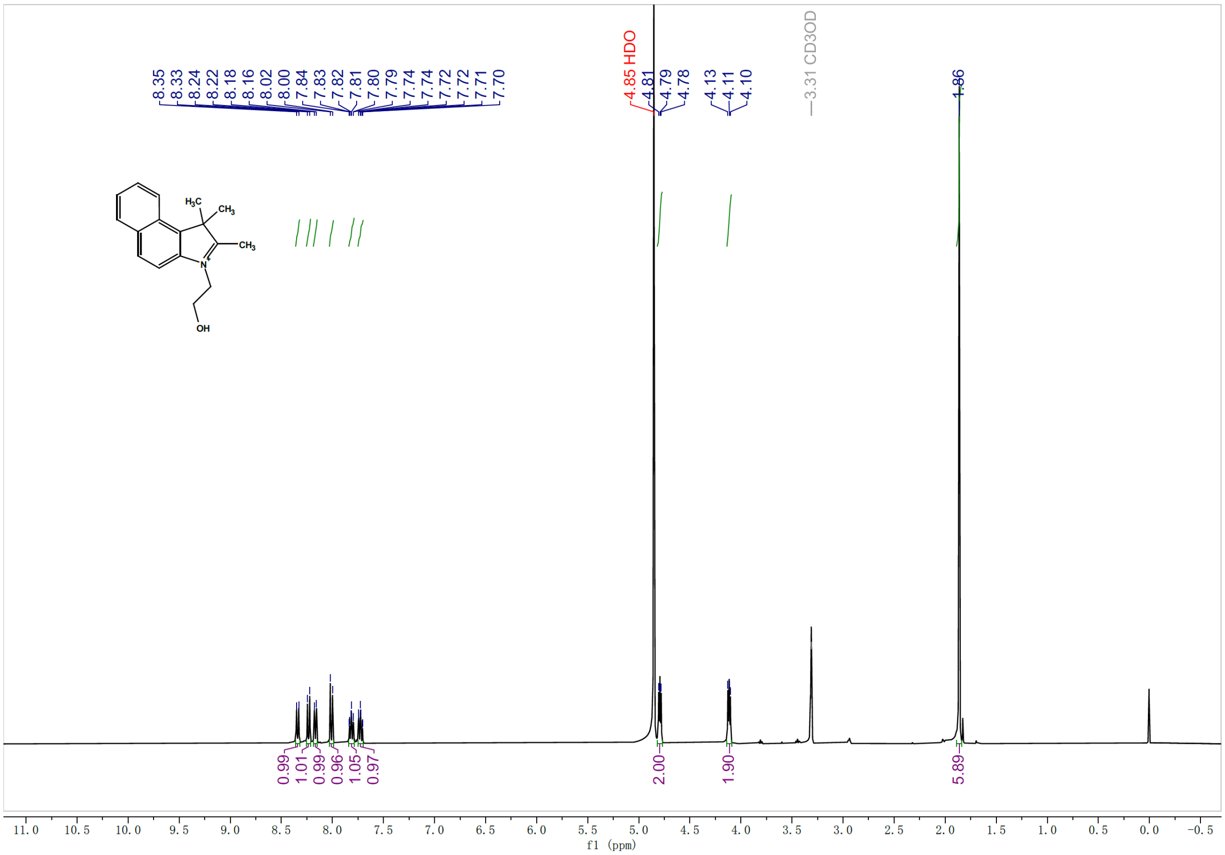


**$^1$H NMR spectrum of compound S2**

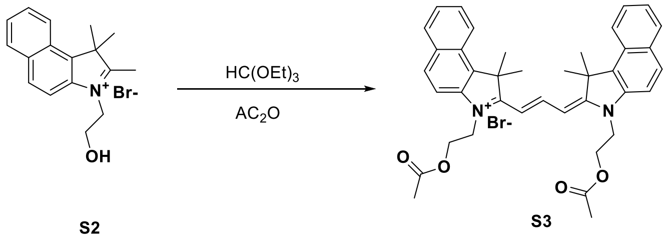


**Compound S3**: A mixture of compound **S3** (450 mg, 1.35 mmol), **triethyl orthoformate** (120 mg, 0.808 mmol), and acetic anhydride (4.5 mL) in a pressure tube was heated at 120°C and stirred for 16 h. The mixture was then cooled to r.t. The solvent was removed under reduced pressure to obtain a residue, which was purified by a silica gel chromatography column, eluting with DCM: MeOH=15:1 to afford the compound (330 mg, 72%) as a purple solid.

$^1$H NMR (400 MHz, Chloroform-*d*) δ 8.69 (t, *J* = 13.3 Hz, 1H), 8.11 (d, *J* = 9.2 Hz, 2H), 7.96-=7.92 (m, 4H), 7.74 (d, *J* = 13.3 Hz, 2H), 7.63 (td, *J* = 7.0, 1.3 Hz, 2H), 7.50 (t, *J* = 8.4 Hz, 2H), 7.45 (d, *J* = 8.9 Hz, 2H) 4.82 – 4.69 (m, 8H), 2.05 (s, 12H), 1.77 (s, 6H).

MS (ESI) calcd for $C_{39}H_{41}N_2O_4^+$ [M]$^+$ 601.31, found 601.17.

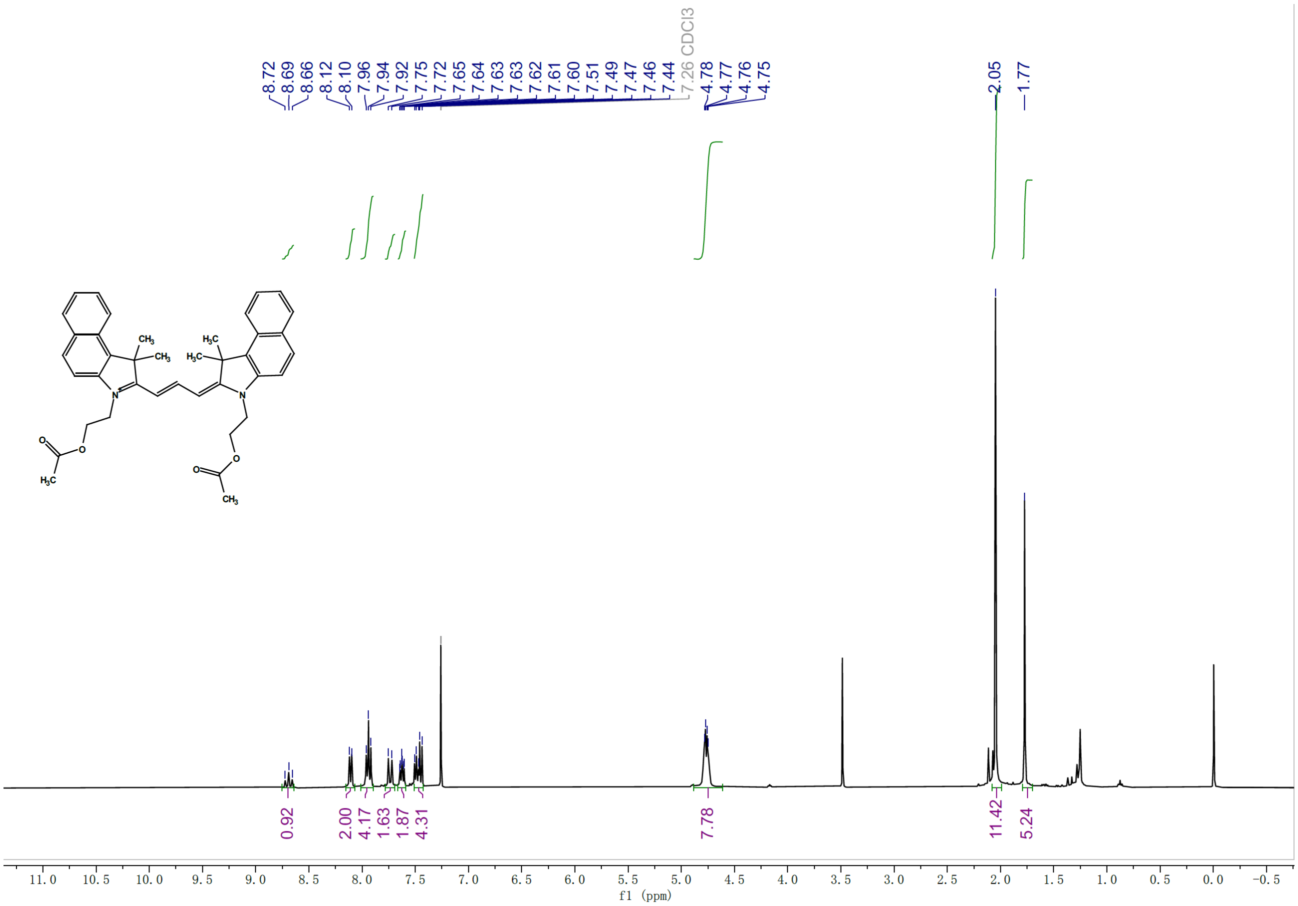


**$^1$H NMR spectrum of compound S3**

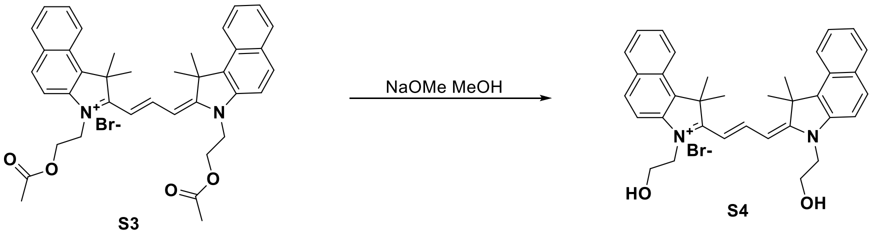


S3 S4

**Compound S4**: To a mixture of compound **S3** (280 mg, 0.411 mmol) in MeOH (6.0 mL) was added NaOMe (111 mg, 2.06 mmol) at r.t. The mixture was stirred for 3 h before concentrated under reduced pressure to obtain a residue, which was purified by a silica gel chromatography column, eluting with DCM: MeOH=30:1 to 10:1 to afford the compound (210 mg, 86%) as a purple solid.

$^1$H NMR (400 MHz, Methanol-*d*$_4$) δ 8.83 (t, *J* = 13.5 Hz, 1H), 8.29 (d, *J* = 8.5 Hz, 2H), 8.03- 7.99 (m, 4H), 7.69-7.65 (m, 4H), 7.51 (t, *J* = 7.5 Hz, 2H), 6.56 (d, *J* = 13.5 Hz, 2H), 4.42 (t, *J* = 5.3 Hz, 4H), 4.06 (t, *J* = 5.1 Hz, 4H), 2.11 (s, 12H).

$^{13}$C NMR (101 MHz, Methanol-*d*$_4$) δ 178.31, 150.87, 141.50, 134.71, 133.58, 131.60, 131.13, 129.26, 128.82, 126.25, 123.36, 112.74, 103.66, 60.31, 52.48, 48.08, 28.00.

MS (ESI) calcd for $C_{35}H_{37}N_2O_2^+$ [M]$^+$ 517.28, found 517.16.

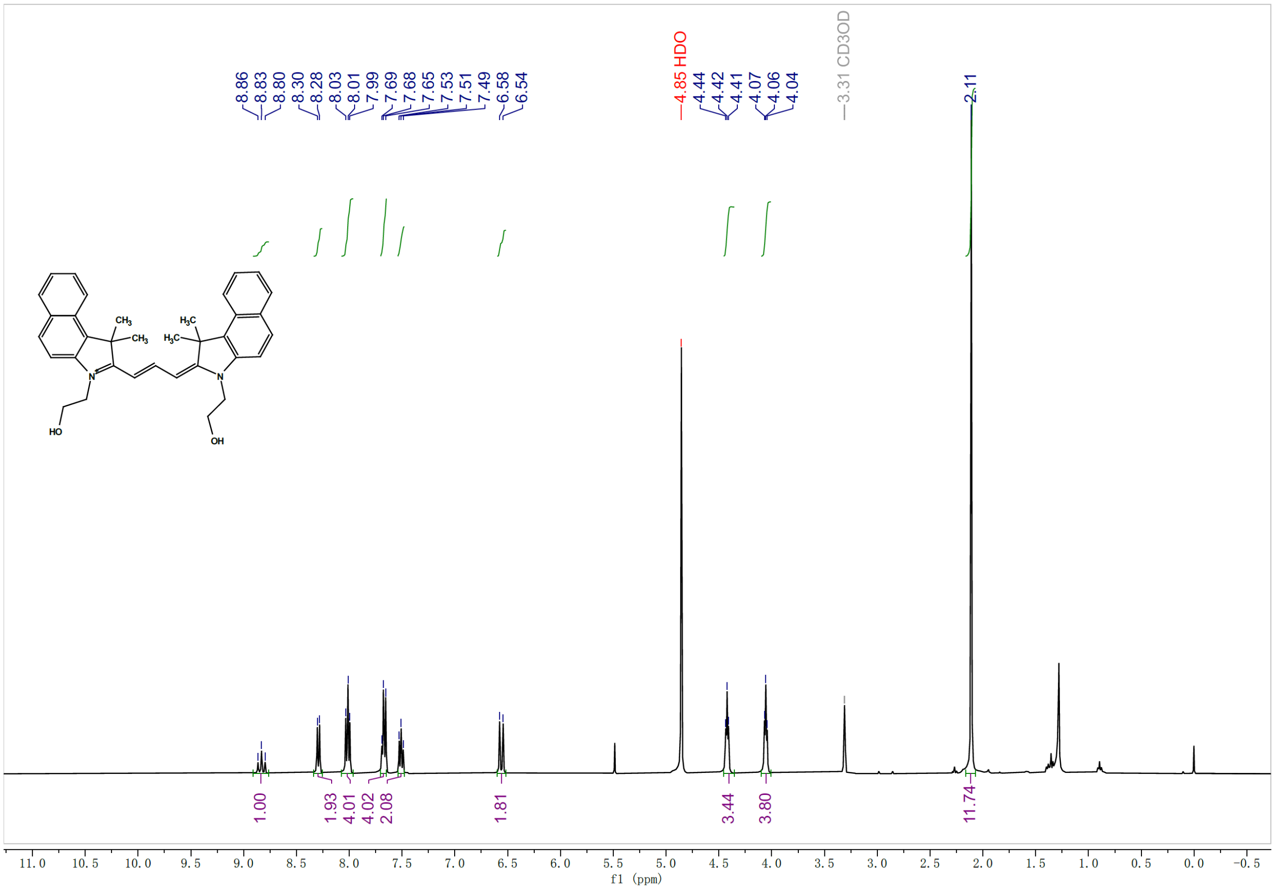


**$^1$H NMR spectrum of compound S4**

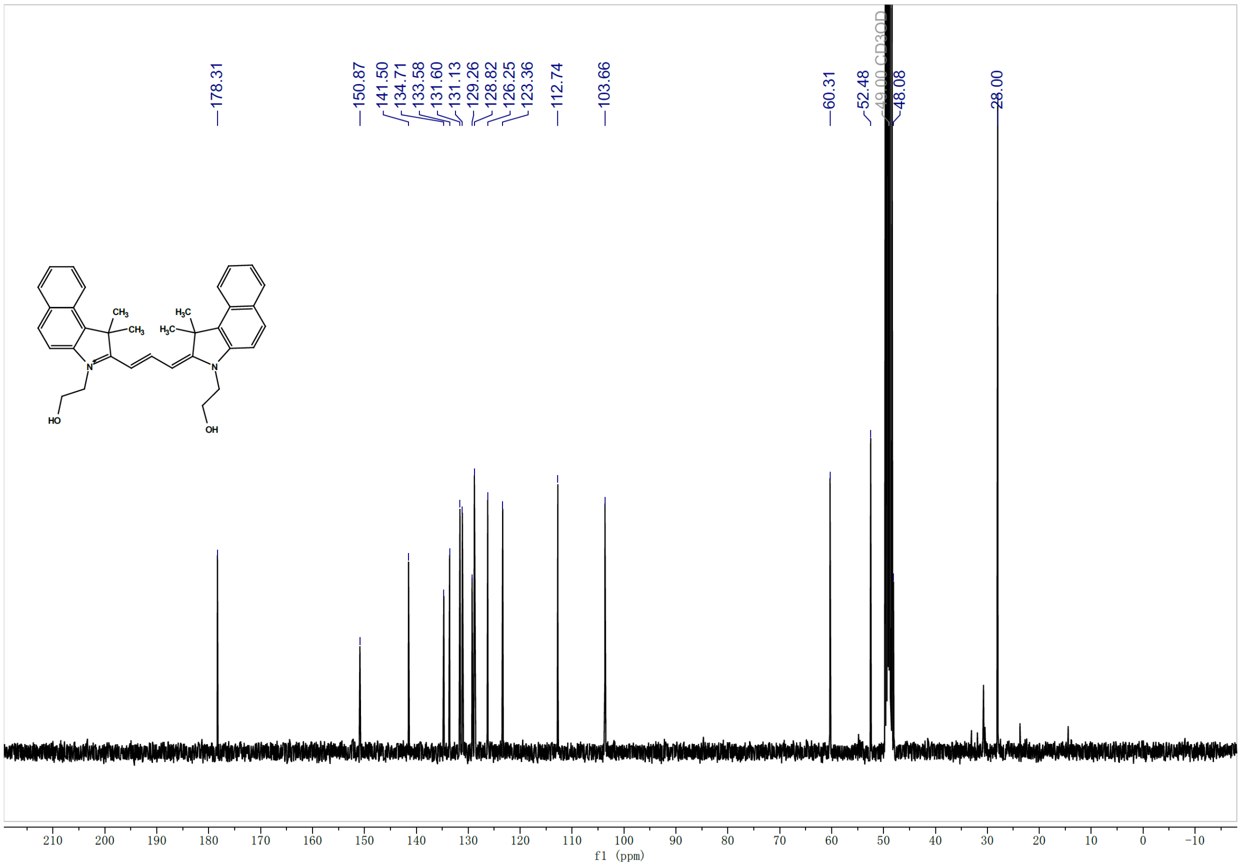


**$^{13}$C NMR spectrum of compound S4**

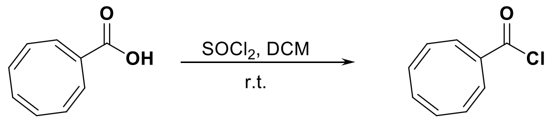

S5

S6

**Compound S6:** To a solution of compound **S5** (60.0 mg, 0.405 mmol) in DCM (5.0 mL) were added $SOCl_2$ (242 mg, 2.03 mmol) and DMF (10 uL) in turn at r.t under nitrogen. The mixture was then stirred for 2 h at r.t. before concentrated under reduced pressure to obtain the crude **S6**, which was used in the next step directly.

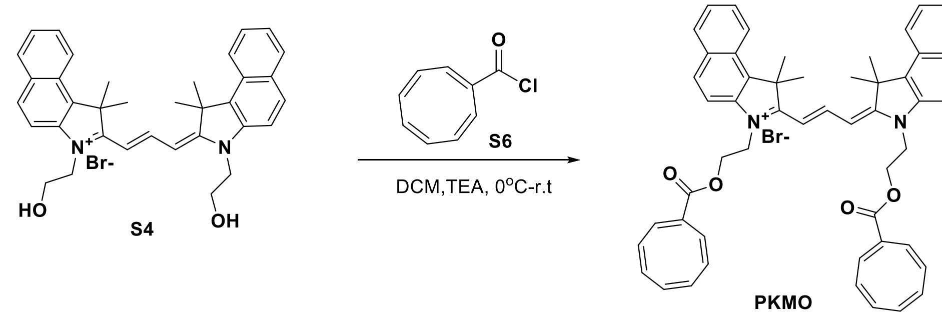


**PKMO:** To a mixture of compound **S4** (35.0 mg, 0.059 mmol) and triethylamine (59.7 mg, 0.590 mmol) in dry DCM (3.0 mL) was added dropwise the solution of compound **S6** (67.5 mg, 0.405 mmol) in dry DCM (1.0 mL) at 0°C under nitrogen. The mixture was stirred at room temperature for 16 h before concentrated under reduced pressure to afford a residue, which was purified by preparative TLC (DCM: MeOH=15:1) to afford crude **PKMO** (40.0 mg, 79%) as a purple solid. The solid was dissolved in DCM (2.0 mL) and added dropwise into diethyl ether (20.0 mL) slowly. The solid was collected and dried under reduced pressure to afford **PKMO** (25.0 mg, 49%) as a purple solid.

$^{1}$H NMR (400 MHz, Methanol-$d_4$) δ 8.78 (t, $J$ = 13.2 Hz, 1H), 8.28 (d, $J$ = 8.5 Hz, 2H), 8.04 - 8.01 (m, 4H), 7.79 – 7.61 (m, 4H), 7.52 (t, $J$ = 7.5 Hz, 2H), 6.82 (s, 2H), 6.64 (dd, $J$ = 13.6, 4.4 Hz, 2H), 5.85 – 5.37 (m, 14H), 4.79 – 4.59 (m, 8H), 2.07 (s, 12H).

$^{13}$C NMR (101 MHz, Methanol-$d_4$) δ 178.13, 166.54, 151.10, 144.86, 140.99, 135.25, 134.71, 134.15, 133.77, 133.66, 132.85, 132.41, 131.91, 131.16, 130.78, 130.01, 129.20, 128.90, 126.39, 123.36, 112.54, 103.97, 62.03, 52.48, 44.45, 28.03.

MS (ESI) calcd for $C_{53}H_{49}N_2O_4^+$ [M]$^+$ 777.37, found 777.28.

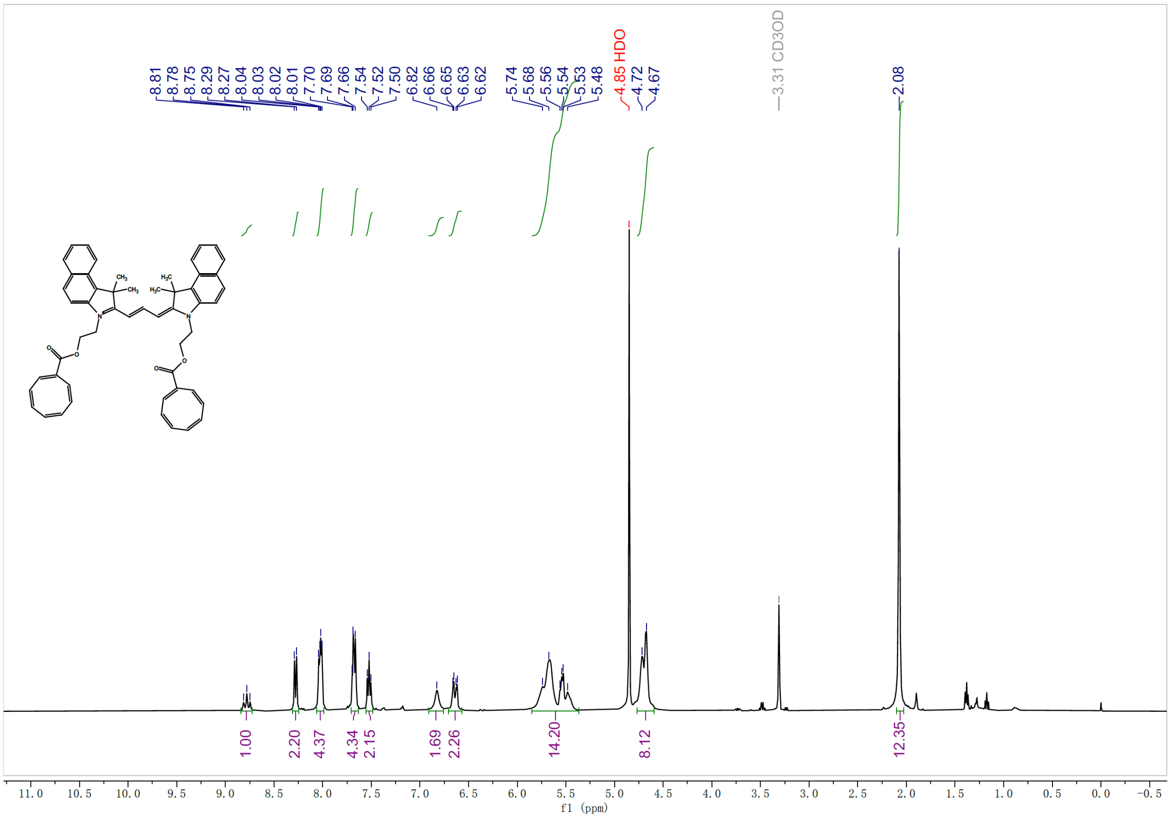


**$^{1}$H NMR spectrum of compound PKMO**

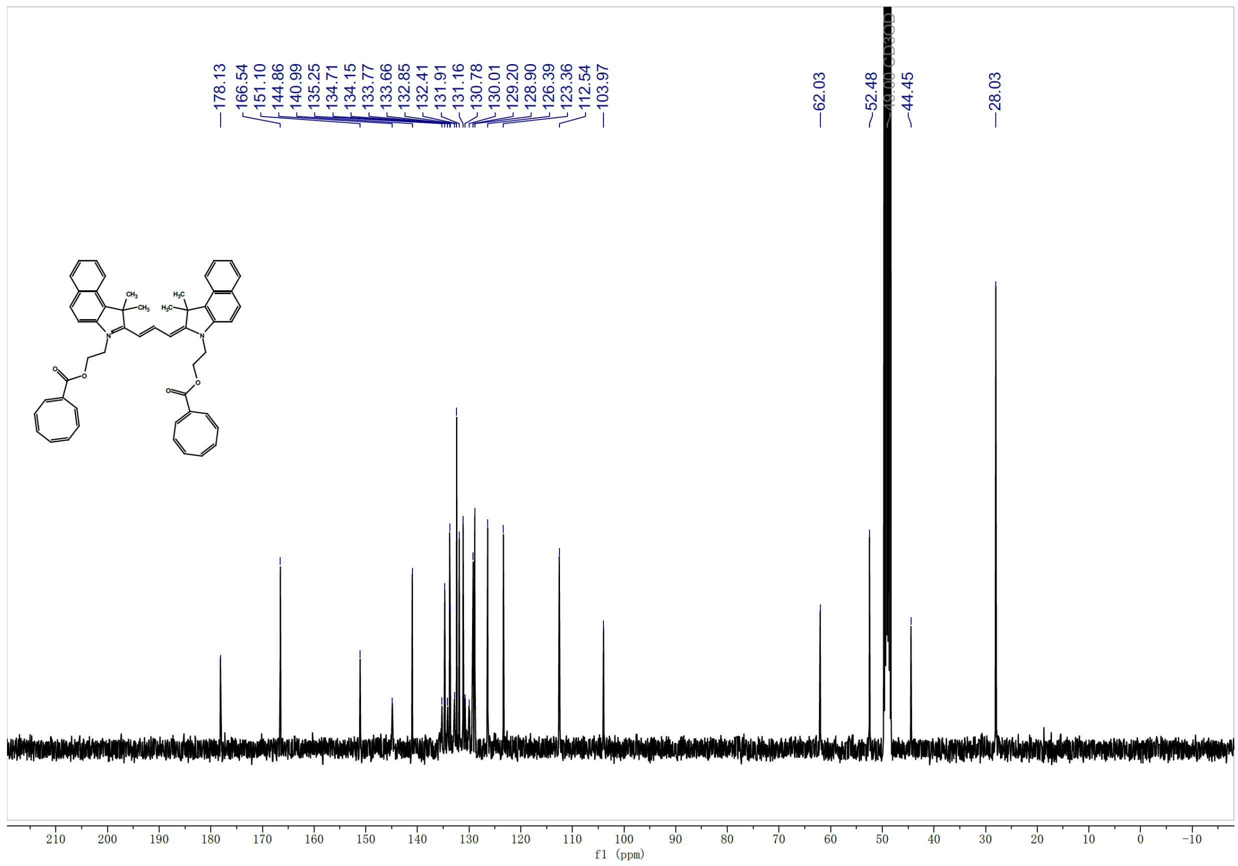


**$^{13}C$ NMR spectrum of compound PKMO**

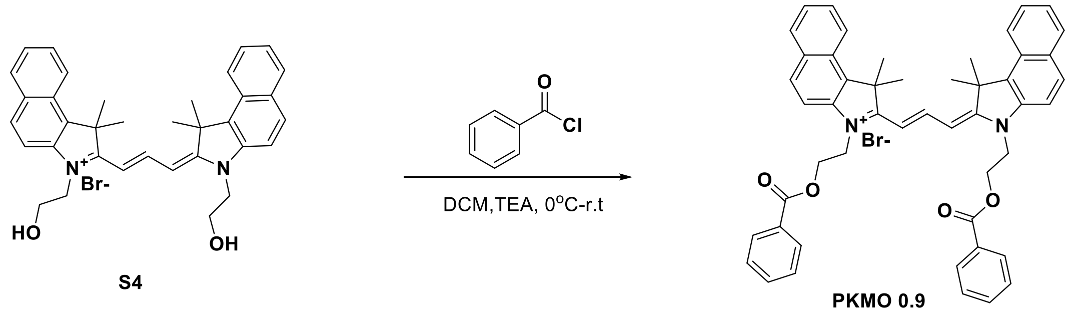


S4 PKMO 0.9

**PKMO 0.9:** To a mixture of compound **S4** (25.5 mg, 0.043 mmol) and triethylamine (21.6 mg, 0.213 mmol) in dry DCM (3.0 mL) was added dropwise the solution of benzoyl chloride (29.8 mg, 0.213 mmol) in dry DCM (0.5 mL) at 0°C under nitrogen. The mixture was stirred at room temperature for 16 h before concentrated under reduced pressure to afford the residue, which was purified by preparative TLC (DCM: MeOH=20:1) to afford crude **PKMO 0.9** (30.0 mg, 97%) as a purple solid. The solid was dissolved in DCM (1.5 mL) and added dropwise into diethyl ether (15 mL) slowly. The solid was collected and dried under reduced pressure to afford the **PKMO 0.9** (20.0 mg, 58%) as a purple solid.

$^1$H NMR (400 MHz, Methanol-$d_4$) δ 8.51 (t, *J* = 13.5 Hz, 1H), 8.21 (d, *J* = 8.6 Hz, 4H), 8.00 - 7.96 (m, 4H), 7.84 (dd, *J* = 8.4, 1.3 Hz, 4H), 7.73 (d, *J* = 8.8 Hz, 4H), 7.65 (t, *J* = 7.7 Hz, 2H), 7.49 (t, *J* = 7.6 Hz, 2H), 7.41 (td, *J* = 7.4, 1.4 Hz, 2H), 7.30 (t, *J* = 7.7 Hz, 4H), 6.52 (d, *J* = 13.4 Hz, 2H), 4.86 (t, *J* = 4.8 Hz, 4H), 4.73 (t, *J* = 5.0 Hz, 4H), 1.92 (s, 12H).

$^{13}$C NMR (101 MHz, Methanol-$d_4$) δ 178.06, 167.49, 150.78, 140.68, 134.70, 134.44, 133.54, 131.75, 131.06, 130.68, 130.40, 129.47, 129.09, 128.85, 126.32, 123.24, 112.30, 103.87, 62.40, 52.32, 44.65, 27.79.

MS (ESI) calcd for $C_{49}H_{45}N_2O_4^+$ [M]$^+$ 725.34, found 725.27.

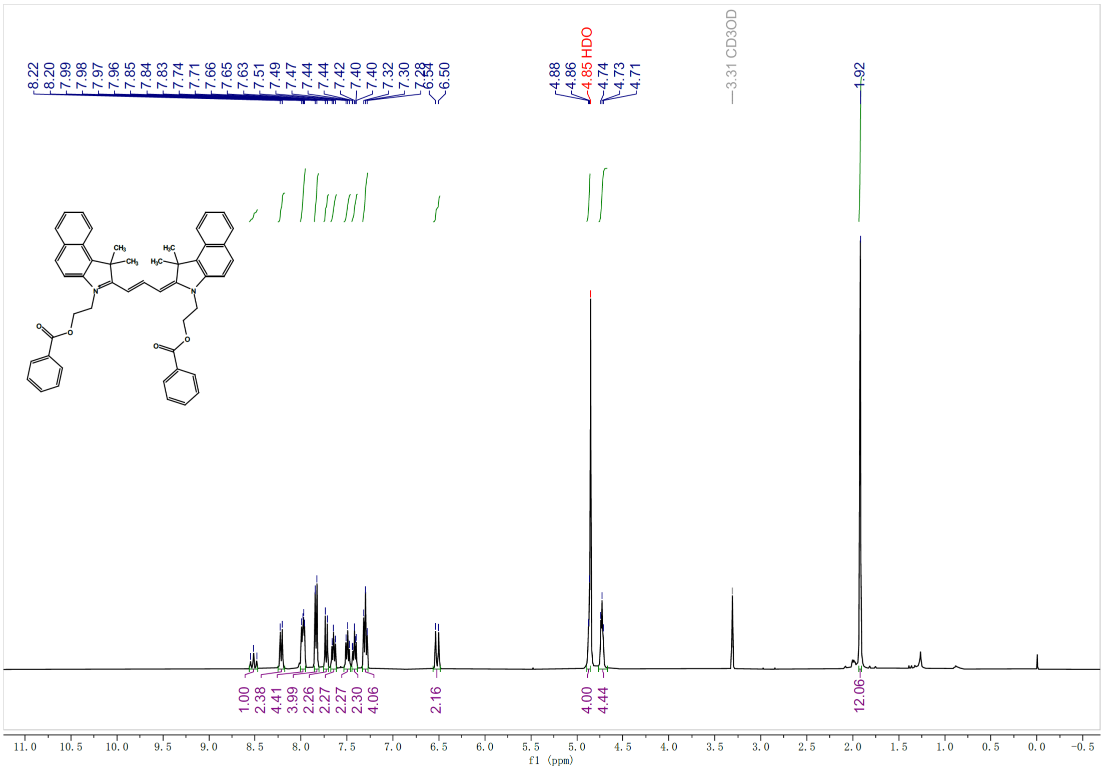


**$^1$H NMR spectrum of compound PKMO 0.9**

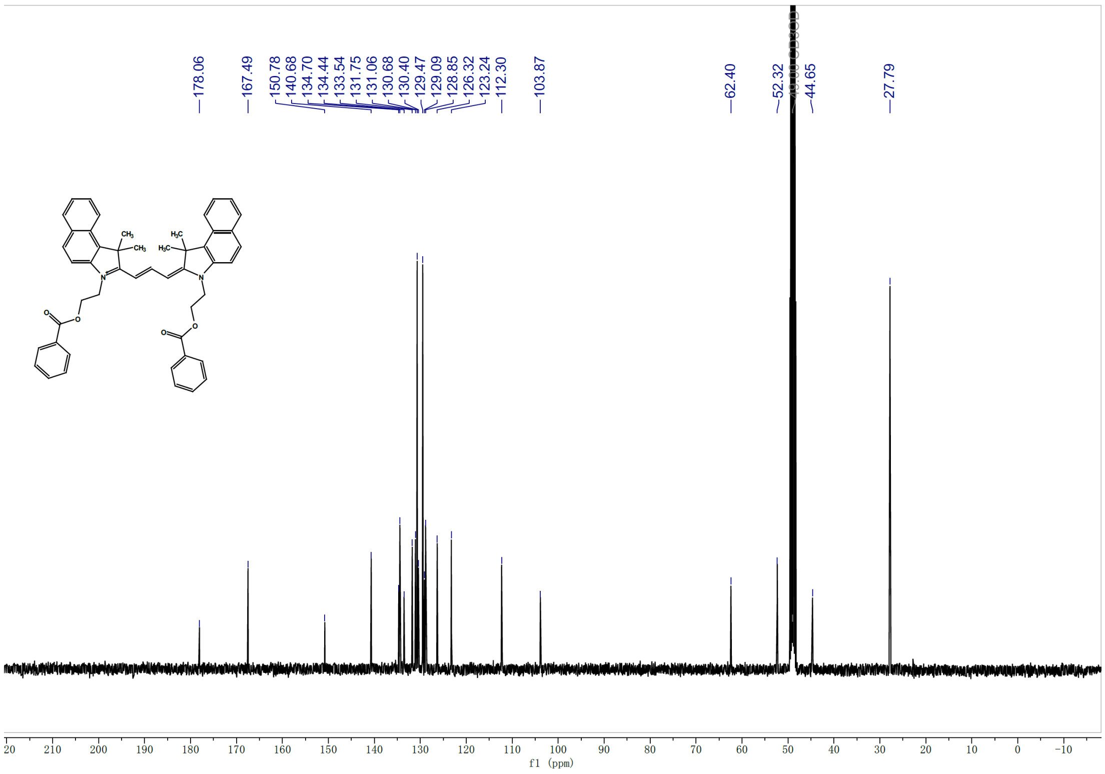


**$^{13}C$ NMR spectrum of compound PKMO 0.9**
